# B7-H3 Orchestrates Fibro-Immune Spatial states in Pancreatic Cancer

**DOI:** 10.64898/2026.11.16.762837

**Authors:** Assya Legrini, Tengyu Zhang, Mari-Claire McGuigan, Colin Wood, Leonor Patricia Schubert Santana, Luke McNickle, Phimmada Hatthakarnkul, Kathryn Pennel, Claire Kennedy Dietrich, Yoana Doncheva, Ghazal Latifi, Josefina Martinez Vasquez, Holly Leslie, Jean Quinn, Colin Nixon, Kai Rakovic, Alexandra S McLaren, Michael Doukas, Joanne Edwards, Nigel B Jamieson

**Affiliations:** School of Cancer Sciences, University of Glasgow, Glasgow, United Kingdom; University Department of Surgery, Glasgow Royal Infirmary, Glasgow, United Kingdom; Histology Department, Cancer Research UK Scotland Institute, Glasgow, United Kingdom; Department of Pathology, Erasmus MC-University Medical Centre Rotterdam, Rotterdam, The Netherlands

## Abstract

Pancreatic ductal adenocarcinoma (PDAC) remains one of the most lethal cancers in the western world and is largely resistant to treatment. A major barrier to effective treatment is the highly desmoplastic, immunosuppressive tumour microenvironment, where the organisation of tumour and stromal cells limit immune infiltration. B7-H3 (*CD276*) is an emerging immune checkpoint marker investigated in phase I and Ib/2 clinical trials in PDAC, yet its biological role within the tumour environment remains poorly understood. We performed an integrated multiscale spatial profiling study to map B7-H3 expression across regional and single-cell spatial contexts in treatment-naïve resected human PDAC. Our results demonstrate that B7-H3 is not simply overexpressed in cancer but may act as a spatial organiser of the PDAC microenvironment, associated with a transition from epithelial-intrinsic tumour architecture to fibro-inflammatory immune states that underpin immune suppression, aggressive disease and poor prognosis.

**Graphical abstract:** 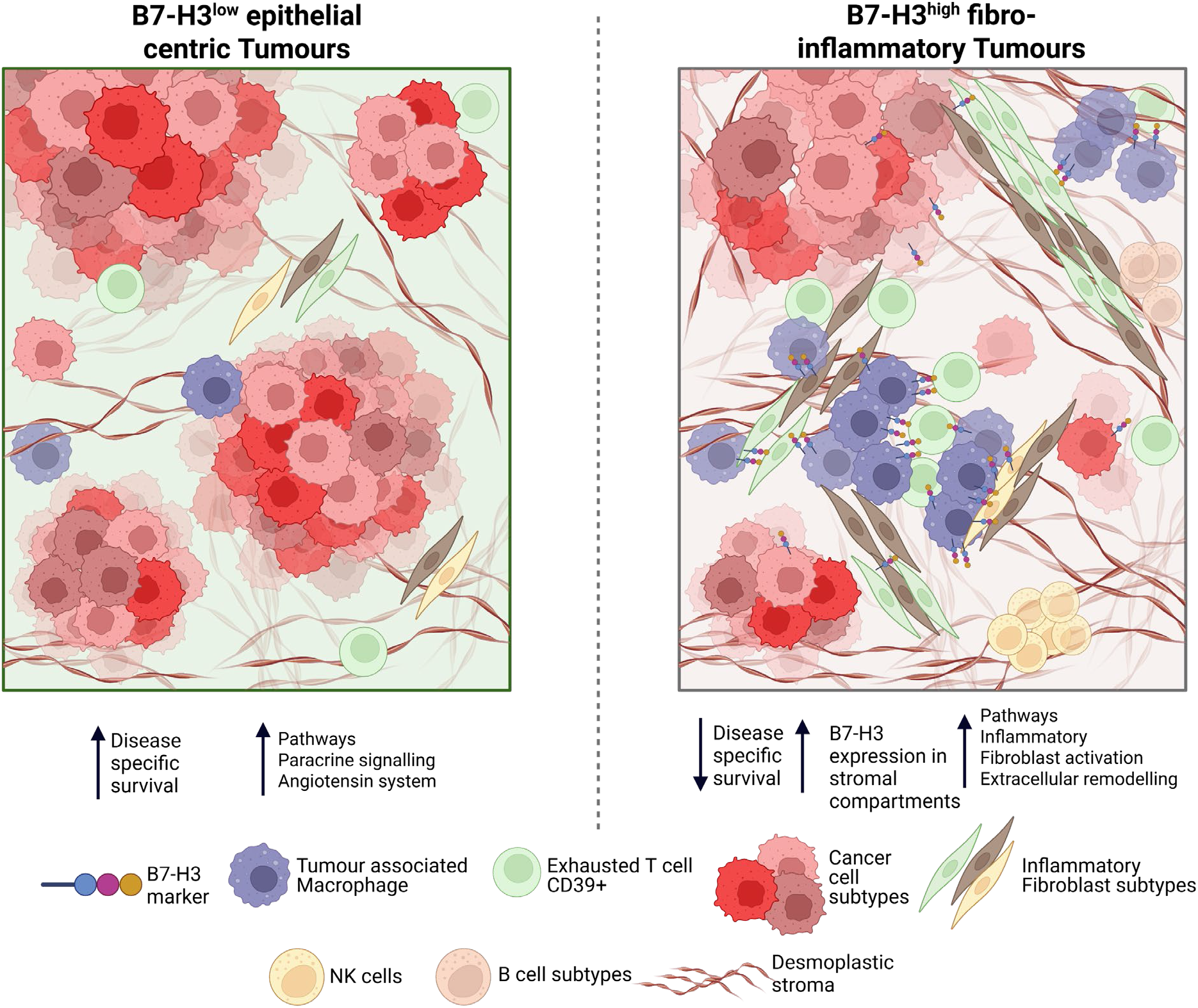

## Introduction

Pancreatic ductal adenocarcinoma (PDAC) remains one of the most lethal malignancies in the western world, with <8% of patients surviving five years (1). Despite research efforts, therapeutic advances have remained limited, with most patients experiencing disease recurrence following surgery and adjuvant chemotherapy (2, 3). However, recent phase III clinical trial, RASolute 302(UK), using KRAS inhibitors have shown promise (4). Therapeutic resistance has made it difficult for successful treatment progression. This is partly attributed to the highly desmoplastic and immunosuppressive tumour microenvironment (TME), which is characterised by a dense fibroblast population and low tumour immune infiltration, resulting in extensive spatial compartmentalisation (5–7).

Immune checkpoint pathways represent attractive therapeutic targets in PDAC, however, clinical responses using monotherapy have been disappointing compared to other solid tumours, partly due to its dense fibrotic TME. This disconnect highlights a critical gap in our understanding of how immune checkpoint molecules are organised within the spatial context of the pancreatic tumour microenvironment, and how their expression relates to the complex interactions between epithelial, stromal, and immune compartments (8–10). Dual treatment with novel immune checkpoint inhibitors may offer a therapeutic solution. Preclinical research demonstrates combining KRAS inhibition with immunotherapy results in tumour regression, with significant restructuring of the immunosuppressive microenvironment and improved T-cell infiltration in PDAC mouse models (11, 12).

B7-H3 (*CD276*) is an immune checkpoint molecule belonging to the immune-regulatory B7 family including PD-L1, implicated in both a co-stimulatory and co-inhibitory immune role, modulating T-cell activation and contributing to immune evasion (13–17). Though this marker is expressed in normal tissue at the mRNA level, it has limited protein expression, suggesting a tightly regulated post-translational mechanism that appears dysregulated within the cancer setting (18). Functionally, B7-H3 is thought to promote tumour growth, immune suppression and metastasis via multiple signalling pathways and is overexpressed in multiple solid tumours (19–21). This dual immunological and tumour-intrinsic role has established B7-H3 as a promising target, with on-going phase III trials in small cell lung cancer and is being evaluated in phase I and Ib/2 clinical trials for pretreated, advanced PDAC patients (22–24). Unlike classical immune checkpoints, the lack of B7-H3 protein expression in normal tissue ensures a highly targeted approach, establishing B7-H3 as an attractive therapeutic target in the pancreatic cancer landscape (25).

Despite rising interest, the phenotypic identity of B7-H3 expressing cells and the spatial interactions through which B7-H3 shapes the microenvironment remain poorly understood. Studies have attributed B7-H3 expression to a variety of cell types including tumour cells, fibroblasts, macrophages, dendritic cells, B and T cells (26–28). Regardless of its phenotypic expression, elevated B7-H3 consistently associates with poor prognosis (19) and immune-suppressive traits including T-cell inhibition and T-cell exhaustion (29, 30). Additionally, B7-H3 remains an orphan ligand, with reported receptor pairs remaining controversial as none have been conclusively validated in human tissue (31–33). It is unclear whether B7-H3 is simply expressed a range of specific cell types, or whether it helps define higher-order spatial tumour states characterised by coordinated epithelial, stromal, and immune interactions. Resolving this distinction is essential for understanding the biological role of B7-H3 within cancer, as well as providing rational for deployment of B7-H3 targeted therapies in PDAC.

This study applied an integrated multiscale spatial profiling strategy to map B7-H3 expression across regional and single-cell spatial contexts in resected treatment-naïve human PDAC. We aimed to define the cellular populations, neighbourhood architectures, and spatial interactions associated with B7-H3 expression, and to determine whether this checkpoint marker stratifies PDAC into distinct states with prognostic relevance. Our findings reveal that B7-H3 defines a fibro-inflammatory tumour state in PDAC, repositioning this checkpoint molecule from a passive biomarker to a potential active organiser of tumour architecture.

## Results

### Study design and cohort overview

We spatially profiled two independent treatment-naive resected PDAC TMA cohorts, named discovery and validation. The discovery PDAC cohort comprised of 28 patients represented by multi-regional cores, with an average 4 cores per patient (Figure 1.a). This cohort was utilised for initial hypothesis data generation using a regional spatial protein assay (GeoMx WTA assay, n = 27) (Figure1.b). The validation PDAC cohort was made up of 79 patients, with one core per patient (Figure 1.a). Serial sections of this cohort were utilised for spatial-omics profiling, undergoing regional spatial transcriptomics (GeoMx WTA assay, n = 61), single cell spatial proteomics (CosMx IO protein assay, n = 74), and single cell spatial transcriptomics (CosMx 6K discovery assay, n = 48) (Figure 1.b).

**Figure 1:**
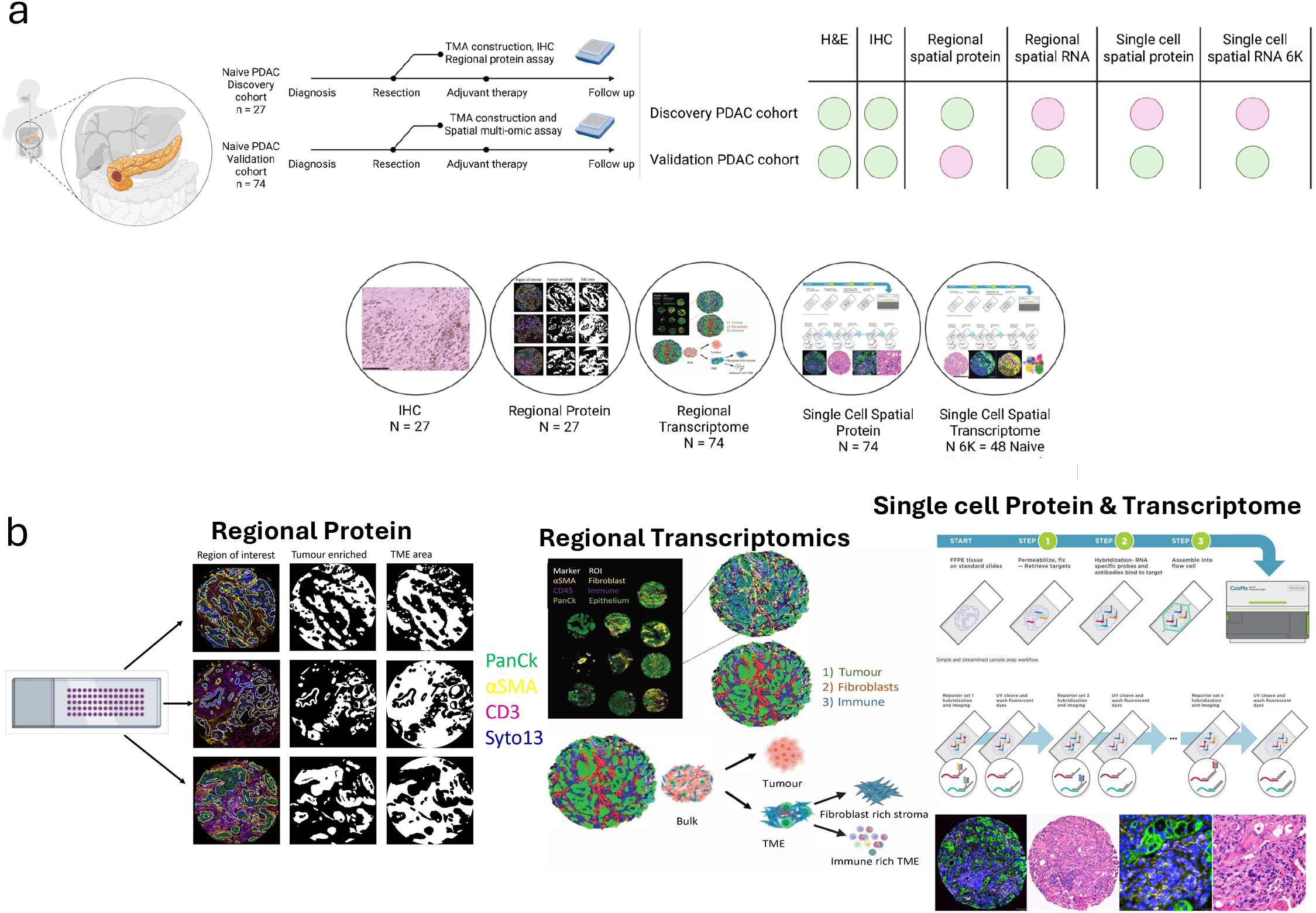
Introduction to Pancreatic Cancer tissue cohort, technology and assays used. **a)**. Schematic showing naïve and neoadjuvant pancreatic cancer cohorts used throughout the study, the table (right) showing the associated assays used, green indicates data is available for the associated assay, red indicates no data available. The circles below indicate the patient number per cohort associated with technology, **b).** Schematic showing types of spatial-omic assays deployed. Regional protein was generated using the GeoMx Human Immune-Oncology (IO) protein assay, samples were fluorescently stained with Syto13, PanCk, CD3 and αSMA, masks, shown in white, were selected for Tumour enriched (PanCk positive) and tumour microenvironment (TME). Regional transcriptomics was generated using the GeoMx Human Whole transcriptome atlas (WTA) assay. Samples were fluorescently stained with Syto13, PanCk, CD45 and αSMA, and masks were selected for Tumour (PanCk positive – green mask), fibroblasts rich stroma (αSMA positive – red mask) and Immune rich TME (CD45 positive – blue mask).Single cell spatial protein and single cell transcriptome were generated using the CosMx Human Immune-Oncology (IO) protein panel and the CosMx Human 6K Discovery panel respectively. All samples were fluorescently stained for DAPI (blue), PanCk (green), CD298/B2M (cyan), CD45 (magenta) and CD68 (yellow).

### B7-H3 associates with regional protein tumour microenvironment compartments and survival in PDAC

Regional spatial proteomic profiling was first performed to characterise the tumour microenvironment (TME) in the discovery treatment-naïve resected PDAC cohort. Strong concordance was observed between regional protein markers and chromogenic IHC for CD3 (R = 0.83, p < 0.001) and CD8 (R = 0.84, p < 0.001), with moderate concordance for CD68 (R = 0.68, p < 0.001) (Figure 2.a-c). Regional protein expression revealed marked differences between epithelial and TME areas. As expected, Pan-Cytokeratin (PanCk) expression was enriched in tumour compartments, whereas immune and stromal markers including CD3, CD4, αSMA and B7-H3 were significantly elevated within TME compartments (Figure 2.d-f), though B7-H3 was also expressed within the tumour (Figure 2.e). B7-H3 expression showed strong positive correlations with the immune and stromal markers FAPα (R=0.69, FDR<0.001), LAG3 (R=0.6, FDR<0.001), CD40 (R=0.56, FDR<0.001), CTLA4 (R=0.51, FDR<0.001), CD25 (R=0.75, FDR<0.001), SMA (R=0.47, FDR<0.001) and CD127 (R=0.75, FDR<0.001) in TME compartments (Figure 2.f and Table 1.b). These associations suggest that B7-H3 is embedded within immune and fibroblast environments.

**Figure 2:**
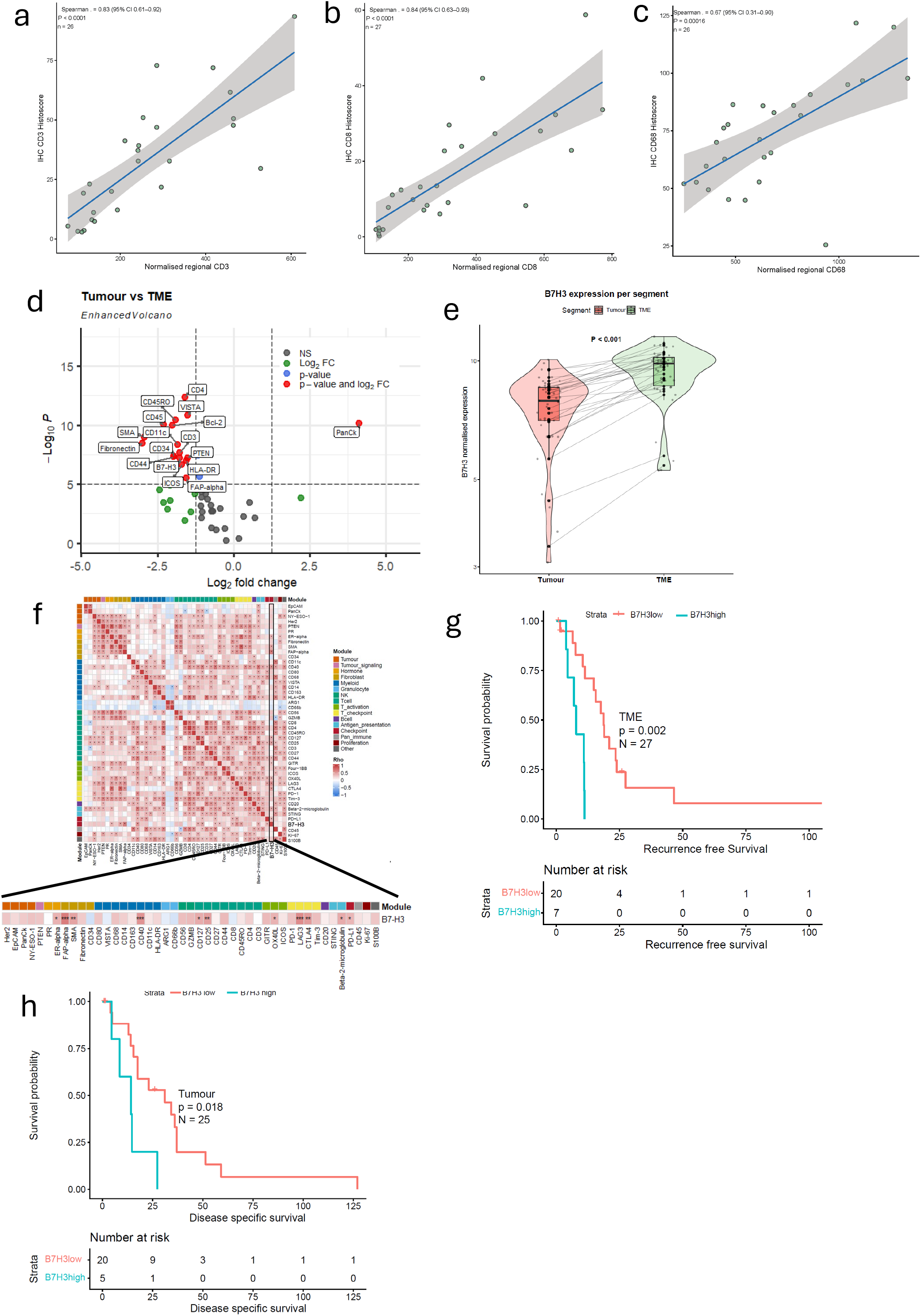
Regional protein profiling defines B7-H3 associated immune and stromal states in PDAC. **a-c).** Regression scatter plots comparing a). CD3, b). CD8, c). CD68 positivity staining in each core, comparing Histo-scores generated by IHC with average normalized regional protein marker expression per patient. Spearman correlation coefficient and p-value presented. Grey shading denotes 95% confidence interval of correlation coefficient. **d).** Volcano plot demonstrating protein marker differential expression levels based on comparison of Tumour versus tumour microenvironment (TME) regions. Dashed line indicates significance thresholds set (fold change=1.25, p value (BH adjusted) =0.05), NS = non-significant, FC = fold change. **e**). Violin Boxplot showing pairwise B7-H3 normalized expression per segment per patient, p value generated from linear mixed-effects model with segment as a fixed effect and a random intercept for patient, accounting for paired inter-patient sampling **f).** Correlation heatmap showing significant markers associated with B7-H3 within the TME, spearman’s correlation test with BH adjusted p values, significant values are indicated by a star. Markers are grouped by module, indicated by different colours. **g-h).** Kaplan Meier curves (Log-Rank test) in ranked B7-H3 expression, stratified by **g).** recurrence free survival in TME compartments, **h).** disease specific survival in Tumour compartments

**Table 1.a.**
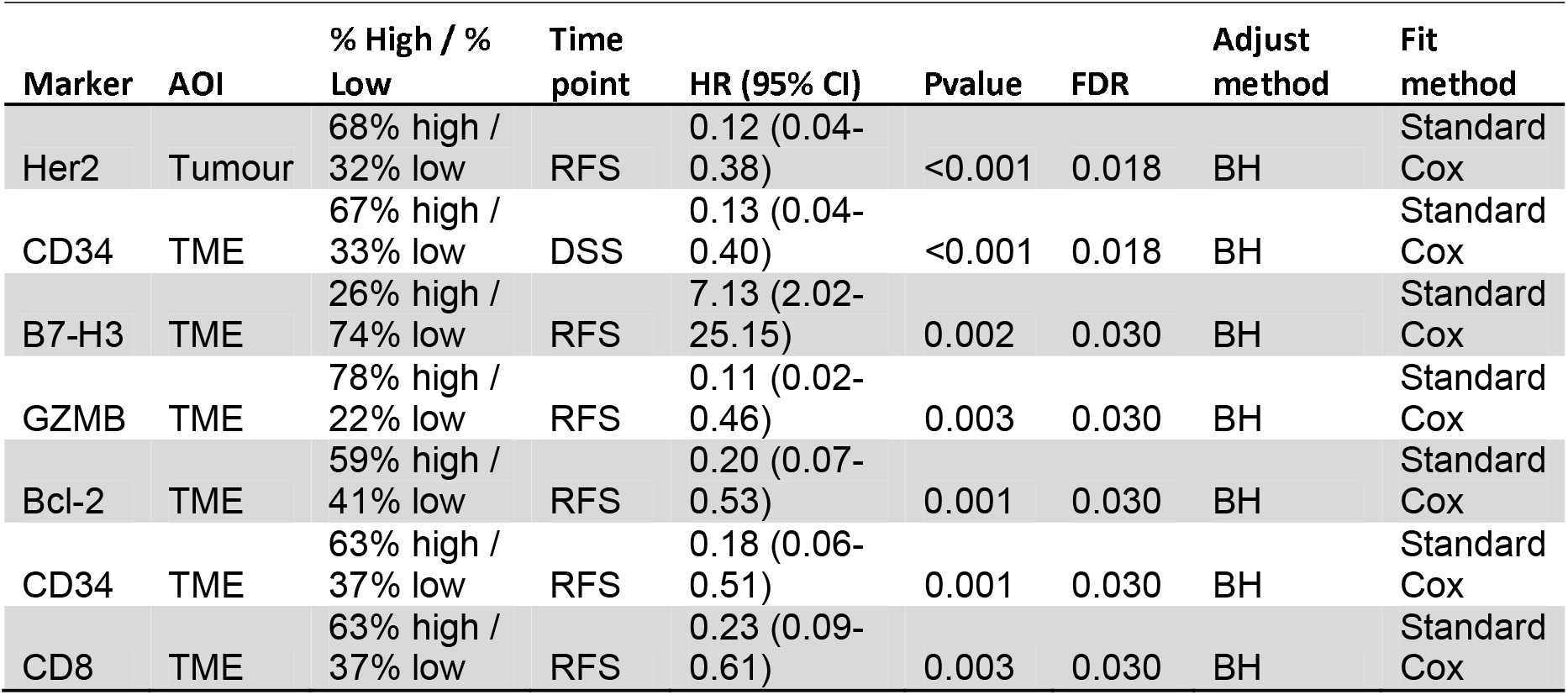
Summary of regional protein biomarker signature density for disease specific survival (DSS) and recurrence free survival (RFS) in tumour and tumour microenvironment (TME) segments in treatment naive pancreatic cancer discovery cohort. Cut-off method established per phenotype and segment generated using maximal selected rank statistics. Log Rank (Mantel-Cox) p value and Univariate cox regression hazard ratio (HR) shown with 95% confidence interval (CI) per marker, per segment, with BH adjusted p values (FDR).

| Marker | AOI | % High / % Low | Time point | HR (95% CI) | Pvalue | FDR | Adjust method | Fit method |
|---|---|---|---|---|---|---|---|---|
| Her2 | Tumour | 68% high / 32% low | RFS | 0.12 (0.04-0.38) | <0.001 | 0.018 | BH | Standard Cox |
| CD34 | TME | 67% high / 33% low | DSS | 0.13 (0.04-0.40) | <0.001 | 0.018 | BH | Standard Cox |
| B7-H3 | TME | 26% high / 74% low | RFS | 7.13 (2.02-25.15) | 0.002 | 0.030 | BH | Standard Cox |
| GZMB | TME | 78% high / 22% low | RFS | 0.11 (0.02-0.46) | 0.003 | 0.030 | BH | Standard Cox |
| Bcl-2 | TME | 59% high / 41% low | RFS | 0.20 (0.07-0.53) | 0.001 | 0.030 | BH | Standard Cox |
| CD34 | TME | 63% high / 37% low | RFS | 0.18 (0.06-0.51) | 0.001 | 0.030 | BH | Standard Cox |
| CD8 | TME | 63% high / 37% low | RFS | 0.23 (0.09-0.61) | 0.003 | 0.030 | BH | Standard Cox |

**Table 1.B.** Top significant regional protein markers correlated with B7-H3 marker in tumour microenvironment segments of treatment naive pancreatic cancer discovery cohort, generated from heatmap in (f), Spearman’s correlation coefficient test used with coefficient (r), BH adjusted p values (FDR) and number of Area’s of illumination (AOIs) shown.

| Marker | Module | r | Pvalue | FDR | AOI number |
|---|---|---|---|---|---|
| FAP-alpha | Fibroblast | 0.669 | 3.790E-07 | 1.710E-05 | 48 |
| LAG3 | T_checkpoint | 0.598 | 1.121E-05 | 2.520E-04 | 48 |
| CD40 | Myeloid | 0.558 | 5.533E-05 | 8.300E-04 | 48 |
| CTLA4 | T_checkpoint | 0.514 | 2.551E-04 | 2.870E-03 | 48 |
| CD25 | Tcell | 0.482 | 6.850E-04 | 6.170E-03 | 48 |
| SMA | Fibroblast | 0.47 | 9.819E-04 | 7.360E-03 | 48 |
| CD127 | Tcell | 0.438 | 2.359E-03 | 1.520E-02 | 48 |
| Beta-2-microglobulin | Antigen_presentation | 0.425 | 3.240E-03 | 1.820E-02 | 48 |
| OX40L | T_activation | 0.399 | 6.025E-03 | 3.010E-02 | 48 |
| ER-alpha | Hormone | 0.381 | 8.909E-03 | 4.010E-02 | 48 |
| PD-L1 | Checkpoint | 0.368 | 1.196E-02 | 4.890E-02 | 48 |

To assess the prognostic relevance of spatially resolved protein expression, survival analysis was performed separately within tumour and TME compartments. TME-specific analysis demonstrated elevated B7-H3 expression correlates with poor recurrence-free survival (Table 1.a). We were specifically interested in the biological difference found within extreme B7-H3 expressors. Based on cut point tests and our biological rationale (Supplemental Methods 5 and Supplemental Methods Table 5), the 80^th^ percentile was used to dichotomise B7-H3. This grouping demonstrated elevated B7-H3 expression in tumour compartments associated with poor disease specific survival and elevated B7-H3 expression in TME compartments associated with poor recurrence free survival respectively (Figure 2.g-h), and combining the two compartments non-significantly replicated this trend (Supplemental Figure 1).

### Regional transcriptomics reveals distinct epithelial and stromal programmes associated with *CD276* (B7-H3) expression

To determine whether B7-H3 expression is associated with distinct transcriptional programmes, we performed regional transcriptomic profiling, using the GeoMx WTA assay, in an independent validation PDAC cohort (Figure 1.a). Given the compartment-specific B7-H3 expression patterns observed in regional proteomics (Figure 2), ROIs were further stratified into tumour, fibroblast-rich (αSMA), and immune-rich (TME) compartments (Figure 1.b).

Principal component analysis demonstrated clear separation between epithelial and fibroblast/immune ROIs, confirming compartment-specific gene expression likely driven by differing cell populations (Figure 3.a and Supplemental Figure 2.a). Patients were ranked by mean whole core *CD267 (*B7-H3) transcriptomic expression and dichotomised at the 80^th^ percentile, as defined in the discovery cohort (Supplemental Methods 5). To keep naming conventions consistent, groups were classified as *B7-H3*^high^ (n = 12), and *B7-H3*^low^ (n = 48). Reduced *CD276* (B7-H3) expression across combined compartments was associated with significantly improved survival, with tumour compartments showing a similar, non-significant, trend (Figure 3.b and Supplemental Figure 2.b). *CD276* (B7-H3) expression was significantly elevated in αSMA regions compared with epithelial compartments (pairwise test, FDR<0.001), recapitulating the protein level observation (Figure 3.c and Figure 2.g).

**Figure 3:**
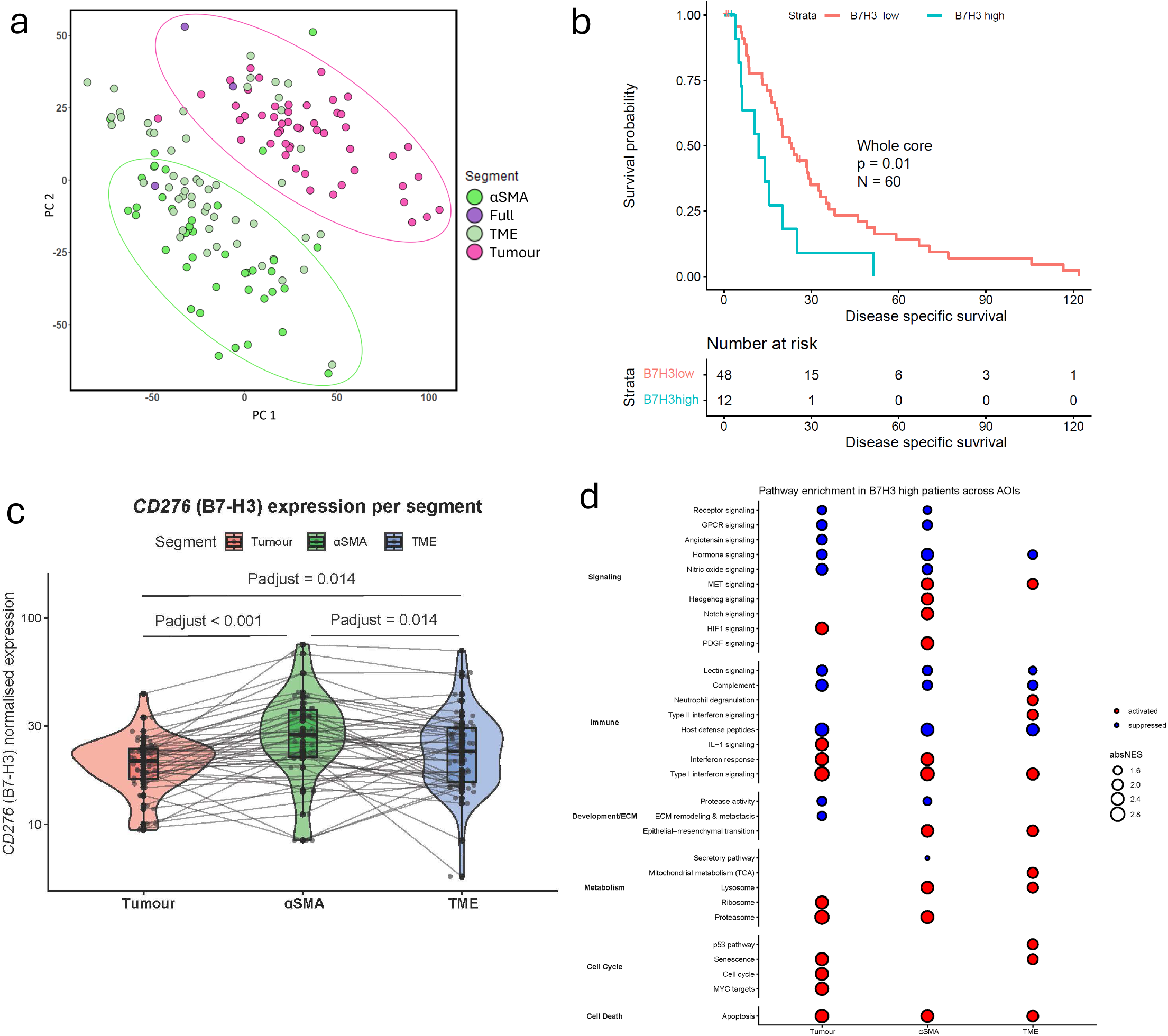
Regional transcriptomic profiling defines B7-H3 high as fibro-inflammatory stromal rich with immune suppression, and B7-H3 low maintains epithelial intrinsic states. **a).** Principle component analysis (PCA) for all segments taken from the validation treatment naïve cohort, annotated and colour indicated by segment. Clusters indicated by ellipses drawn, **b).** Kaplan Meier curves (Log-Rank test) stratified by disease specific survival and in ranked B7-H3 expression across PanCk positive tumour epithelium. Ranked expression generated from 80^th^ percentile as indicated in supplemental methods 5, **c).** Violin Boxplot showing pairwise B7-H3 normalized expression per segment per patient, p value generated from linear mixed-effects model with segment as a fixed effect and a random intercept for patient, accounting for paired inter-patient sampling, **d).** Dot plot showing top significant enriched pathways in B7-H3high segments, pathways used are from Nanostring specific GSEA pathways. Colour of dot indicates activation state, red = activated, blue = suppressed, and size of dot indicates the absolute normalized expression score (absNES) per pathway. Pathways split into overall biological module (far left annotation).

#### Compartment specific analysis helps define CD276 (B7-H3) associated pathways

Pathway analysis comparing *CD267* (B7-H3) ranked patient groups (80^th^ percentile) revealed that *B7-H3*^low^ tumour regions were characterised by significant activation of epithelial signalling programmes, including nitric oxide and angiotensin system signalling (Figure 3.d and Supplemental Table 1). In contrast, *B7-H3*^high^ tumour regions predominantly showed suppression of epithelial programmes alongside relative enrichment of inflammatory signalling pathways, including type I interferon and IL-1 (Figure 3.d). Notably, *B7-H3*^high^ tumour regions did not show enrichment of alternative epithelial-specific signalling programmes, suggesting that the dominant associated biology is not epithelial-intrinsic but immune related. Similarly, *B7-H3*^high^ αSMA and TME regions demonstrated activation of interferon-driven immune programmes, fibroblast activation pathway PDGF, tissue remodelling processes (EMT, MET and Hedgehog), and inflammatory signalling cascades including neutrophil degranulation, and Type I and II interferon response pathways, whereas *B7-H3*^low^ patients were enriched for host defence, nitric oxide and hormone-related signalling pathways (Figure 3.f-g and Supplemental Table 1).Spatial immune cell deconvolution further revealed significantly increased fibroblast (Wilcoxon test, p<0.001) and macrophage (Wilcoxon test, p=0.031) populations in *B7-H3*^high^ TME regions (Supplemental Figure 2.c). These results indicate that *CD267* (B7-H3) expression stratifies PDAC into two transcriptionally and broadly spatial distinct states. The *B7-H3*^low^ phenotype is defined by epithelial-intrinsic signalling programmes with limited immune and stromal related pathways, and the *B7-H3*^high^ phenotype is dominated by fibro-inflammatory, stromal activation and limited epithelial-specific signalling.

### Single cell spatial proteomics demonstrates B7-H3 consistently stratifies PDAC into two reproducible states

#### Iterative cell typing in naïve PDAC using single-cell spatial proteomics

To define the cell types associated with B7-H3 expression, we performed single-cell spatial proteomic profiling (64 plex CosMx IO protein panel) in a serial section of the validation PDAC cohort (Figure 1.a-b). A hierarchical cell-typing strategy integrating protein expression profiles, multiplex immunofluorescence images, and histopathological validation was applied (Figure 4.a), resolving cellular identities into lineages (Figure 4.b), parent cell types (Figure 4.c), and subtypes (Figure 4.d), with subtypes likely reflecting cell states (Supplemental Figure 3.b and Supplemental Table 2).

**Figure 4:**
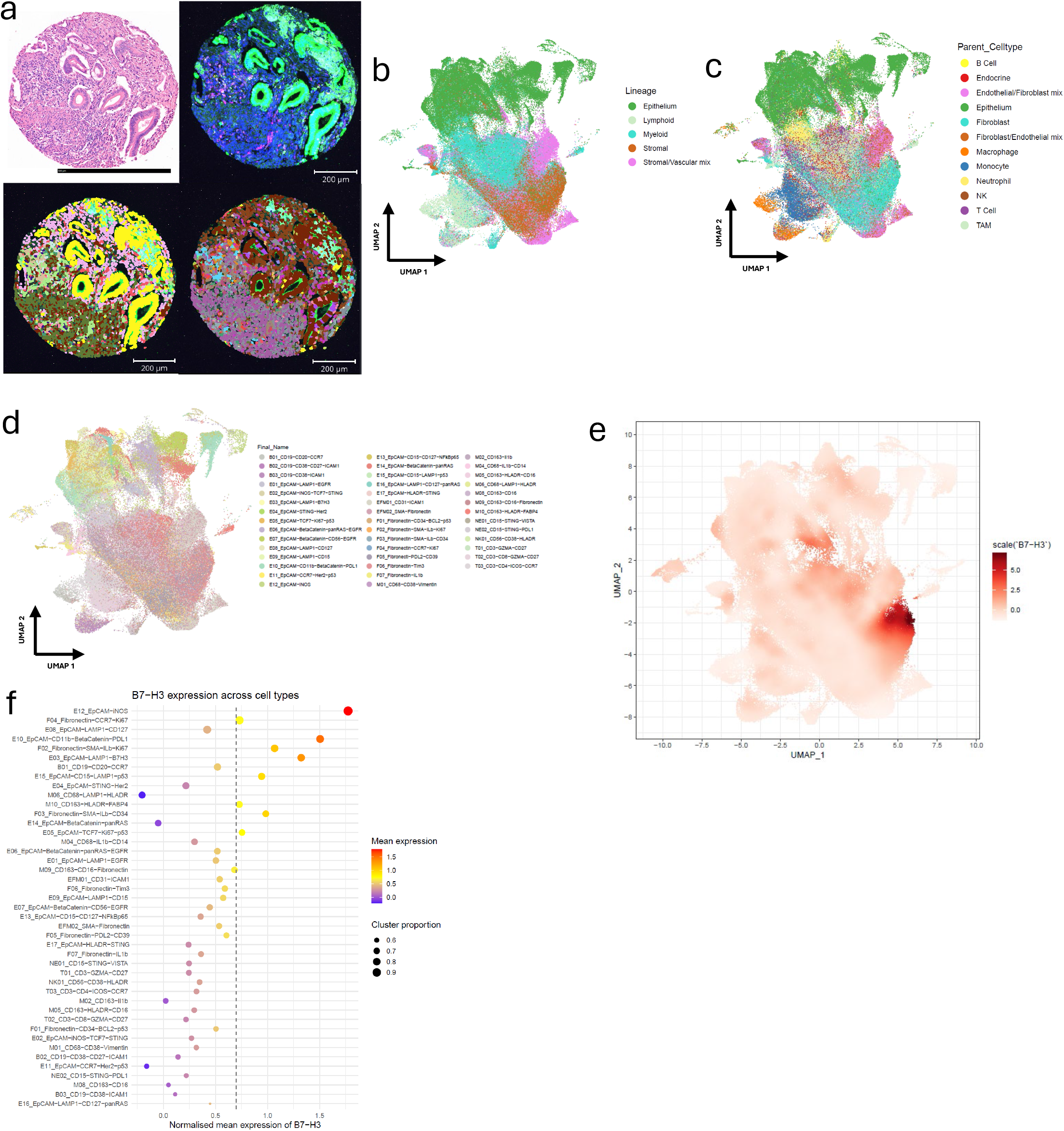
Cell typing reveals B7-H3 enriched epithelial, fibroblast, and myeloid populations in treatment naïve PDAC. **a).** Example core visualisation of HCE (top left), multiplex immunofluorescence (mIF) image stained with DAPI – blue, PanCk – green, CD45 – magenta, CD298/B2M - cyan (top right), overlay of parent cell types on mIF image (bottom left), overlay of fine-grained subclusters on mIF image (bottom right). Bottom 2 are coloured by their respective cell types. **b-d**). Uniform Manifold Approximation and Projection (UMAP) plots representing the hierarchical waterfall cell typing approach utilised, **b).** UMAP showing the overall lineage cell types, **c).** UMAP showing parent cell types, **d).** UMAP showing all data points with final subclusters names. Coloured by cell types indicated in key. **e).** Smoothened B7-H3 expression UMAP on all data points, showing location of enriched protein expression, dark red indicates elevated expression. **f).** Mean B7-H3 expression and cluster proportion of cells expressing B7-H3 per final subcluster, dashed line indicates threshold set.

#### B7-H3 expression is concentrated in specific cellular subtypes

B7-H3 expression was preferential enriched within stromal lineages, and to a lesser extent epithelial and myeloid lineages (Figure 4.e). While B7-H3 expression was relatively uniform across parent cell types, clear differences emerged at the subtype level (Figure 4.f and Supplemental Figure 3.d). Specific subtypes demonstrated elevated B7-H3 expression including epithelial (E03, E05, E10, E12, E15), fibroblast (F02, F03, F04), and macrophages (M10) (Figure 4.f and Table 2). Notably, B7-H3 enriched epithelial subtypes indicated functional states associated with activated signalling and stress-response pathways (Ki67 in E05, iNOS in E12, and LAMP1 and p53 in E15), whereas B7-H3 enriched fibroblast and macrophage populations were characterised by antigen presentation, proliferative and inflammatory signalling features (ILβ and Ki67 in F02, ILβ and CD34 in F03, and HLA-DR and FABP4 in M10) (Table 2).

**Table 2.**
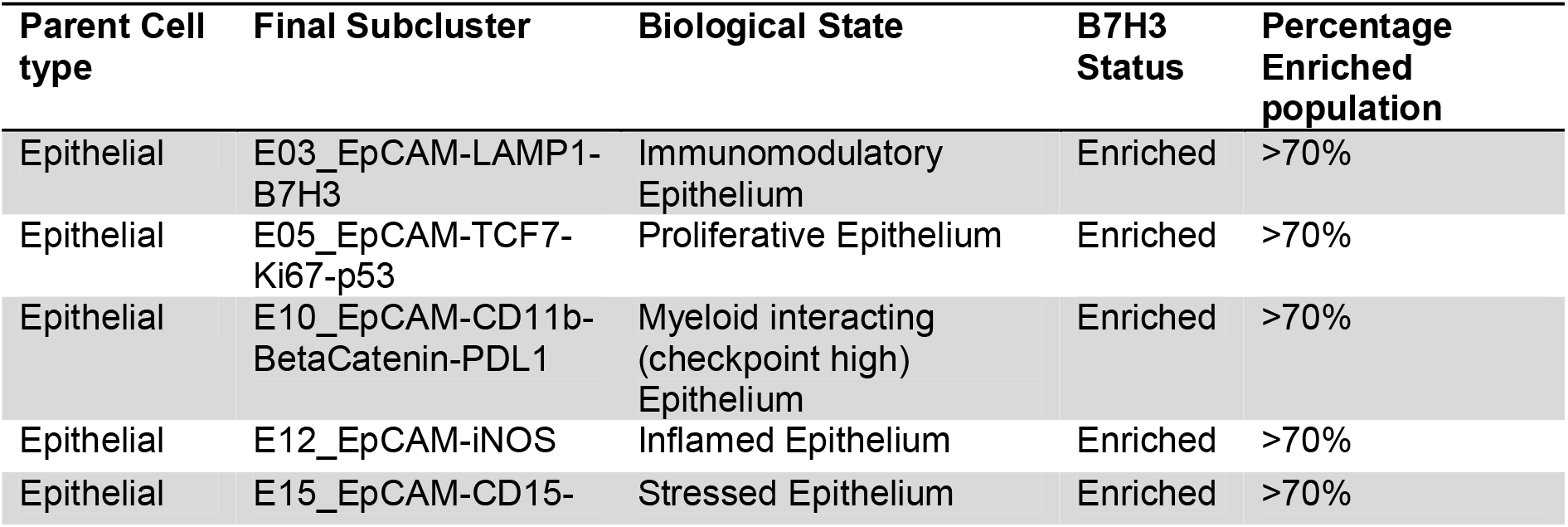
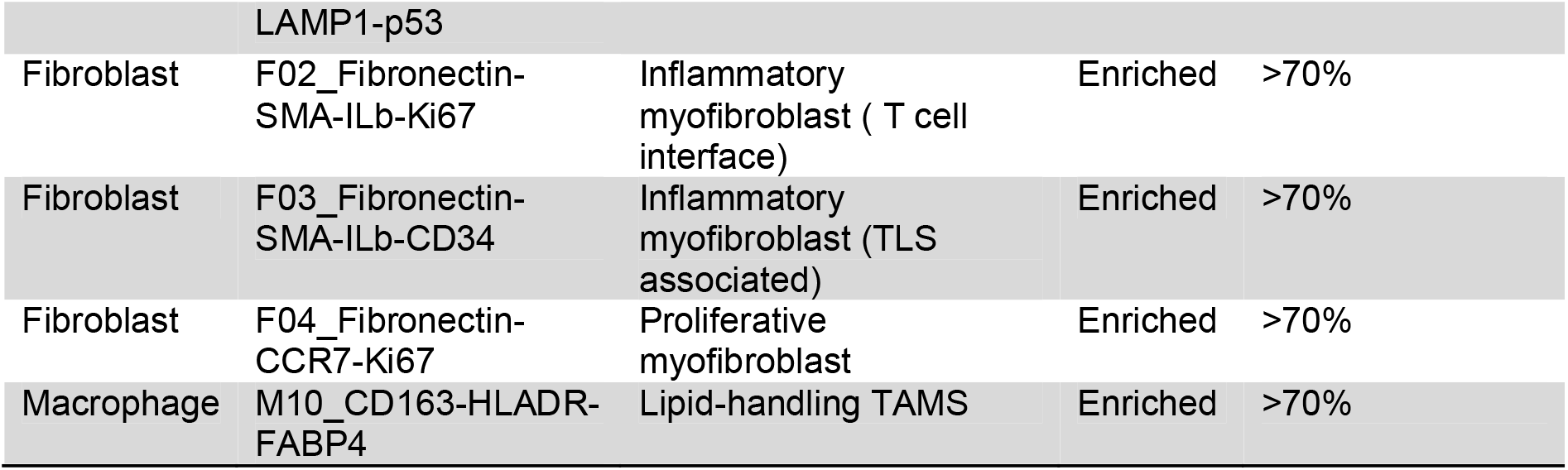
List of top phenotypes globally enriched with B7-H3 marker within treatment naïve pancreatic cancer validation cohort. Parent cell type, final subcluster name and assumed biological state indicated.

#### B7-H3 defines multi-scale tumour microenvironment states in PDAC, characterising fibro-immune spatial architecture in B7-H3^high^ disease while preserving epithelial-centric architecture in B7-H3^low^ disease

To determine whether B7-H3 expression is associated with specific spatial architectures, we performed multi-scale analysis using four complimentary approaches; 1). Micro-neighbourhoods (expression-based clustering network, representing local microenvironments), 2). Macro-niches (cell-type composition, radius-based structure, representing tissue architecture, 3). Spatial domain (aggregated parent cell types, used in gradient based analysis characterising broad microenvironment changes from a central point), and 4). Spatial interaction networks (distance-based differential expression of cell-cell pairings, indicating cellular interaction).

Recurrent micro-neighbourhoods composed of mixed epithelial, stromal, and immune populations were observed, demonstrating that naïve PDAC architecture is organised around multicellular spatial assemblies rather than isolated cell types (Figure 5.a-b). Dominant micro-neighbourhoods were consistently enriched for fibroblasts, tumour-associated macrophages (TAMs), and T cells, defining fibro-immune structures as a core feature of the PDAC microenvironment (Figure 5.b and Supplemental Figure 4.a). B7-H3 expression was most enriched, albeit non-significantly, within a subset of mixed, immune-heavy enriched micro-neighbourhoods (top 3: 25, 7 and 27) (Figure 5.c and Supplemental Figure 4.b). Combined B7-H3 associated micro-neighbourhood groups were generated according B7-H3 enrichment per neighbourhood with a minimum cell count of 200, resulting in B7-H3 enriched micro-neighbourhoods (combination of 25, 7, 27, 29, 5 and 9) demonstrating significant increased expression of fibroblast and neutrophil parent cell types, and reduction of epithelium and B cells compared to B7-H3 depleted micro-neighbourhood (combination of 32, 24, 28, 21, 23 and 20 (Figure 5.d), with heterogenous subtypes observed (Figure 5.e). These results indicate that B7-H3 is not uniformly distributed but instead associated with specific spatial micro-neighbourhoods.

**Figure 5:**
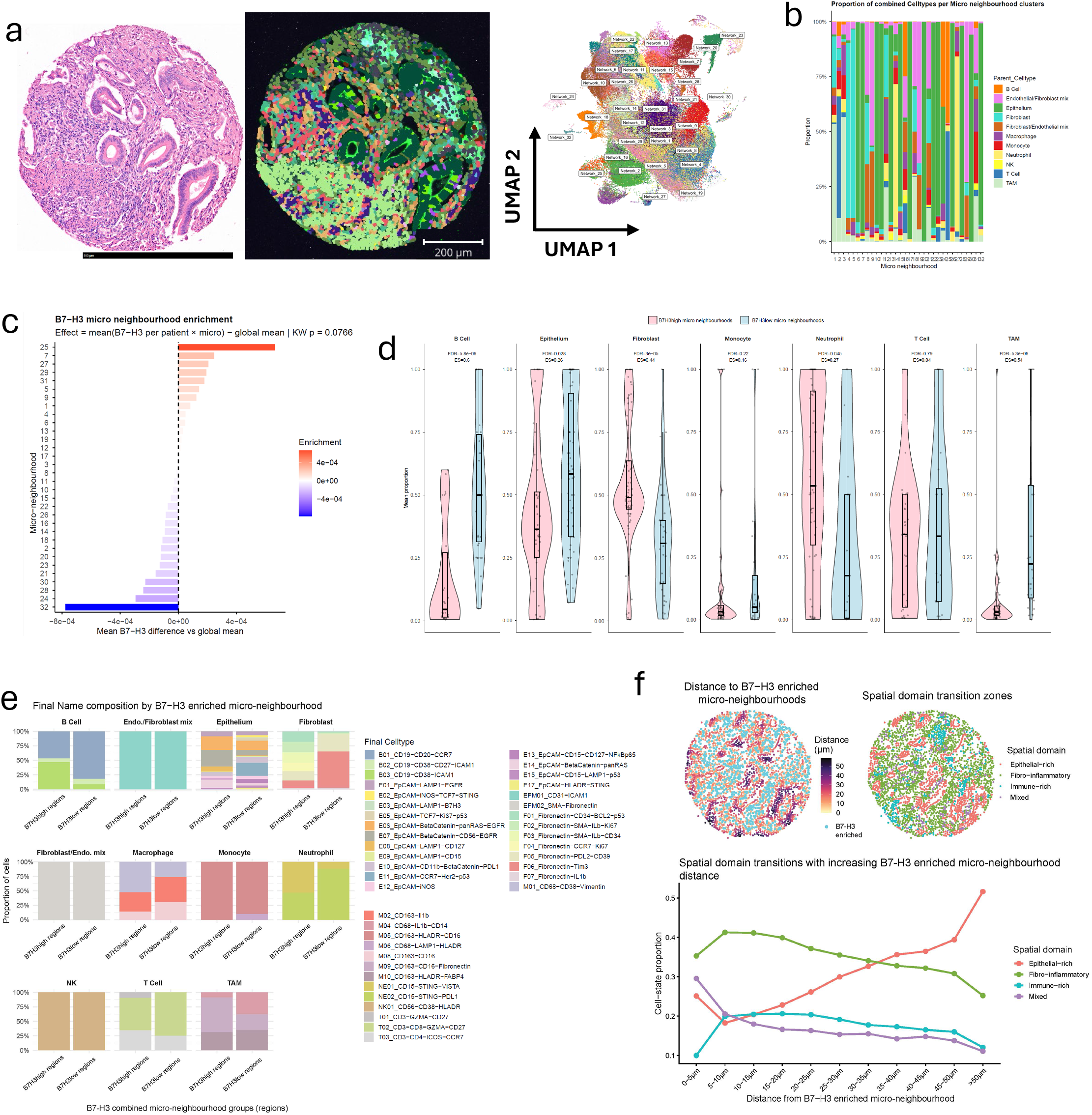
B7-H3 defines multiscale tumour states in PDAC, organising fibro-immune spatial architecture in B7-H3 high disease while preserving epithelial-centric architecture in B7-H3 low disease. **a).** Example core visualisation of HCE (left), overlay of expression micro neighbourhoods on mIF image coloured by neighbourhood cluster (middle), UMAP of micro neighbourhood clusters(right). **b).** Stacked bar chart showing proportion of parent cell types per micro neighbourhood, coloured by cell type. **c).** Ranked bar chart showing mean B7-H3 expression enrichment across micro neighbourhood. **d).** Violin box plot showing mean proportion of parent cell types across B7-H3 combined micro neighbourhoods, selected by enrichment and a minimum of n = 200 cells, resulting in B7-H3 high micro-neighbourhoods classified as combination of 25, 7, 27, 29, 5 and 9, and B7-H3 low micro-neighbourhoods classified as combination of 32, 24, 28, 21, 23 and 20. **e).** Stacked bar chart showing proportion of final subclusters per ranked B7-H3 combined micro neighbourhood, plots are split by parent cell type. **f).** Representative spatial maps from a B7-H3 high PDAC core demonstrating ecosystem organisation relative to B7-H3 enriched regions. Left, spatial distance map showing the Euclidean distance of individual cells from B7-H3 enriched regions (defined by combined micro neighbourhood (25, 7, 27, 29, 5, and 9). Right, classification of local spatial domains based on aggregated parent cell types, including epithelial-rich (epithelial parent cell type), fibro-inflammatory (fibroblasts, Macrophage and TAMs parent cell types), immune-rich (T cell, B cell, monocyte, neutrophil NK parent cell types), and mixed regions (all other cell types). Lower panel, proportion of spatial domain by grouped increasing distance (5μ m increments) from combined B7-H3 high micro neighbourhoods.

Distance analysis demonstrated fibro-inflammatory spatial domains (generated from combined fibroblast, macrophage and TAM parent cell types), were enriched closer to B7-H3 enriched micro-neighbourhoods, peaking at distances 5-10µm (Figure 5.f). Comparatively, epithelial-rich domains progressively increased with greater distance, peaking at distance >50µm (Figure 5.f), indicating a gradient transition from fibro-immune microenvironments toward epithelial-dominant tissue architecture. At the tissue scale, integration of B7-H3 micro-neighbourhood states with macro-niche architecture demonstrated that combined B7-H3 high micro-neighbourhoods preferentially localise within fibro-immune rich macro niches (3 and 7), whereas epithelial-centric niches were depleted (Supplemental Figure 4.c-e). Together, these analyses demonstrate that B7-H3 stratifies PDAC into distinct spatial tumour architecture. B7-H3^high^ tumours are characterised by fibro-inflammatory architectures integrating stromal and immune compartments, whereas B7-H3^low^ tumours retain epithelial-centric spatial organisation. These findings indicate B7-H3 as a spatially relevant focal point of the PDAC tumour microenvironment.

### Point pattern analyses reveal B7-H3 centred fibro-immune clustering

B7-H3 expression per patient was dichotomised at the 80^th^ percentile, as defined in the discovery cohort (Supplemental Methods 5) and used for downstream analysis. Univariate results demonstrated cell type-specific clustering above complete spatial randomness across biologically relevant radius (10-50 µm), with distinct clustering patterns observed between B7-H3 expression states (Figure 6.a). B7-H3^low^ tumours exhibited strong clustering of epithelial subtypes with activation states E02, E04 and E03 (mean degree of clustering range 0.05-0.15), indicating spatial consolidation of epithelial programmes within tumour regions (Figure 6.a and Table 3; Supplemental Figure 5.a and Supplemental Table 2). In contrast, B7-H3^high^ tumours showed preferential clustering of innate immune and inflammatory myeloid populations NK01, M04, B02 and F07 (mean degree of clustering range 0.05-0.07), consistent with fibro-inflammatory cellular states defined above (Figure 6.a and Table 3; Supplemental Figure 5.a and Supplemental Table 2). These findings reinforce that B7-H3 expression is associated with a shift from epithelial-centred spatial organisation to fibro-immune clustering within the PDAC microenvironment.

**Table 3.** Univariate Ripley’s K analysis of cell types clustering across B7-H3 ranked groups. Mean deltaK values are shown for corresponding phenotypes, marker column, per B7-H3 ranked group (B7H3_rank). Positive values indicate clustering above that of expected under spatial randomness. Mean deltaK was calculated across the number of radii indicated in column n_radii

| B7H3_rank | Marker | Mean DeltaK exact | n_radii |
|---|---|---|---|
| High (top 20%) | N01_CD56 | 0.073 | 5 |
| High (top 20%) | NK01_CD56-CD38-HLADR | 0.070 | 5 |
| High (top 20%) | M04_CD68-IL1b-CD14 | 0.059 | 5 |
| High (top 20%) | B02_CD19-CD38-CD27-ICAM1 | 0.057 | 5 |
| High (top 20%) | B01_CD19-CD20-CCR7 | 0.055 | 5 |
| High (top 20%) | F07_Fibronectin-IL1b | 0.054 | 5 |
| High (top 20%) | E12_EpCAM-iNOS | 0.054 | 5 |
| High (top 20%) | E07_EpCAM-BetaCatenin-CD56-EGFR | 0.033 | 5 |
| High (top 20%) | E16_EpCAM-LAMP1-CD127-panRAS | 0.033 | 5 |
| High (top 20%) | M02_CD163-Il1b | 0.031 | 5 |
| High (top 20%) | T01_CD3-GZMA-CD27 | 0.031 | 5 |
| High (top 20%) | B03_CD19_CD38_ICAM1 | 0.029 | 5 |
| Low (bottom 80%) | E02_EpCAM-iNOS-TCF7-STING | 0.150 | 5 |
| Low (bottom 80%) | E04_EpCAM-STING-Her2 | 0.108 | 5 |
| Low (bottom 80%) | N01_CD56 | 0.093 | 5 |
| Low (bottom 80%) | E03_EpCAM-LAMP1-B7H3 | 0.063 | 5 |
| Low (bottom 80%) | F07_Fibronectin-IL1b | 0.062 | 5 |
| Low (bottom 80%) | B01_CD19-CD20-CCR7 | 0.060 | 5 |
| Low (bottom 80%) | M04_CD68-IL1b-CD14 | 0.058 | 5 |
| Low (bottom 80%) | E15_EpCAM-CD15-LAMP1-p53 | 0.057 | 5 |
| Low (bottom 80%) | E08_EpCAM-LAMP1-CD127 | 0.054 | 5 |
| Low (bottom 80%) | B02_CD19-CD38-CD27-ICAM1 | 0.048 | 5 |
| Low (bottom 80%) | NK01_CD56-CD38-HLADR | 0.045 | 5 |
| Low (bottom 80%) | E11_EpCAM-CCR7-Her2-p53 | 0.045 | 5 |

**Figure 6:**
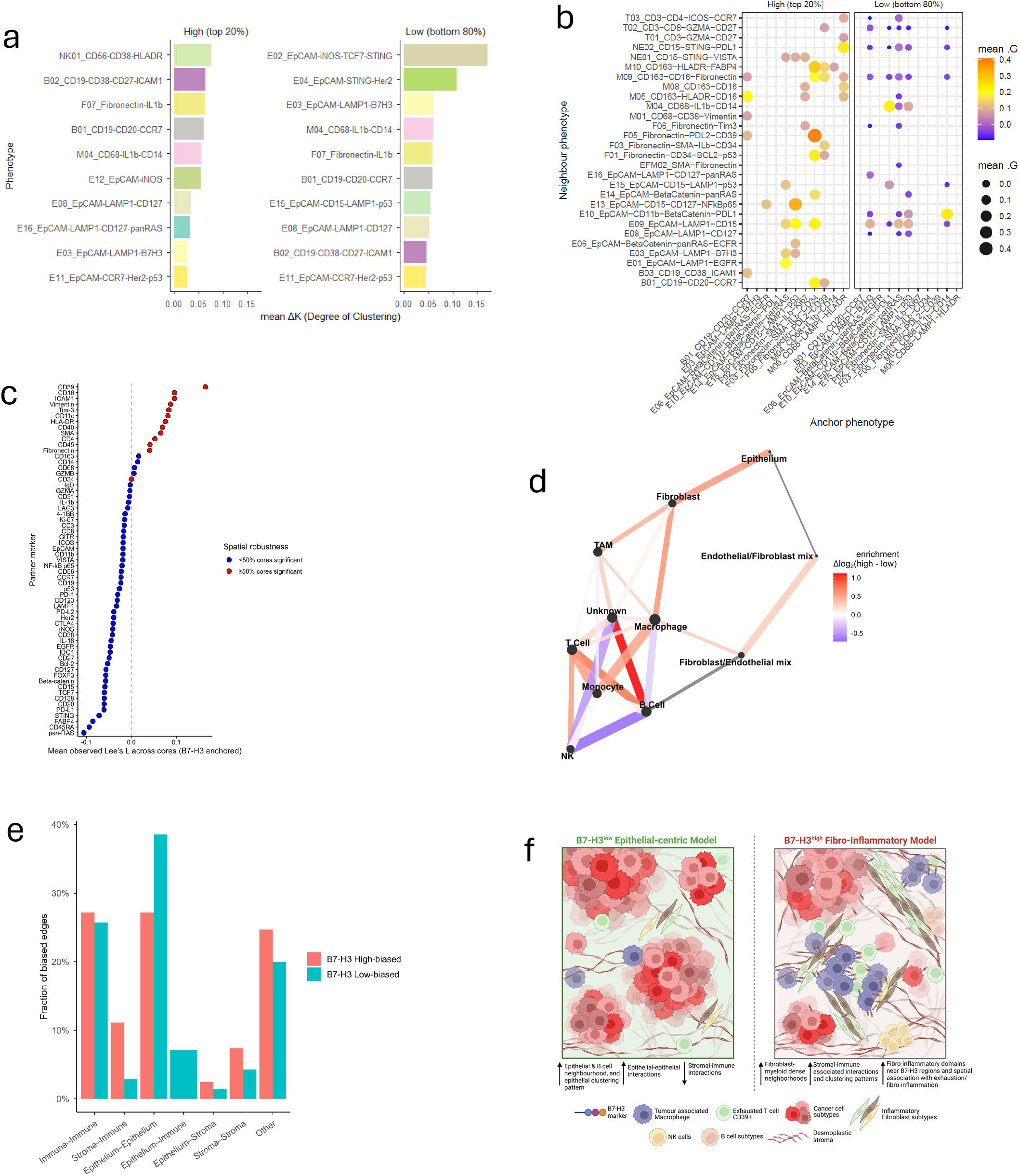
B7-H3 expression defines a spatial reprogramming of the PDAC tumour microenvironment, establishing B7-H3 as a spatial organiser of tumour states. **a).** Ranked bar chart showing Univariate Ripley’s K mean degree of clustering of top fine-grained subclusters split across B7-H3 ranked expression, coloured by cell type. Mean delta K degree of clustering = represents the difference between the observed spatial pattern and the theoretical random distribution. Observed K = the actual count, Exact CSR = complete spatial randomness. **b).** Dot plot showing Bivariate Nearest neighbour mean proximity enrichment between B7-H3 ranked patients and all other final subclusters, coloured by cell types, size of dot indicates enrichment, radius investigated 10-50μm. **c).** Ranked dot plot showing mean observed Lee’s L across cores (range = -0.1 to 0.2), with B7-H3 as the anchor and all other markers as the partner. Spatial robustness is displayed, showing cores with <50% significant coloured blue, and cores with >50% significant coloured red. **d).** Differential enrichment was estimated by aggregating final subcluster-resolved spatial network interactions into parent-level edges, producing effect sizes that reflect both magnitude (enrichment level indicated by colour) and multi-edge (strength of the interaction = line thickness) **e).** Bar chart showing fraction of biased edges aggregated by spatial interaction class. Bars are coloured by B7-H3 ranked expression, red = B7-H3high ranked patients, blue = B7-H3low ranked patients. **f).** Model figure demonstrating the two B7-H3 associated states

#### Spatial mapping reinforces B7-H3^high^ fibro-immune architectures and correlates to immune exhaustion

Bivariate nearest-neighbour analysis revealed non-random spatial proximity between B7-H3 ranked patient groups. At the patient level, B7-H3^high^ tumours exhibited enriched proximity between fibroblast subsets, inflammatory macrophages, and activated lymphocytes, consistent with formation of fibro-inflammatory immune-regulatory niches including F03-F05, F03-M10 and M06-M07 pairs (Figure 6.b and Table 4.a). In contrast, B7-H3^low^ tumours demonstrated limited fibro-myeloid nearest neighbour pairs, supporting a spatial architecture centred on tumour epithelium rather than stromal-immune integration (Figure 6.b and Table 4.a). These results demonstrate B7-H3 expression is associated with short-range coupling of fibroblast and immune populations, reinforcing the concept of B7-H3 centred fibro-immune niches.

**Table 4.a.** Top ranked nearest neighbour cellular interactions across B7-H3 ranks. Top 10 non-self-interactions for each B7-H3 expression group. Anchor denotes the reference cell phenotype and counted denotes the neighbouring cell phenotype. Mean deltaG represents the strength of the nearest-neighbour interaction, averaged across five radii (n_radii = 5). Positive values indicate increased association between named phenotypes.

| **B7H3_rank** | **Anchor** | **Counted** | **Mean deltaG** | **n_radii** |
|---|---|---|---|---|
| High (top 20%) | B02_CD19-CD38-CD27-ICAM1 | T02_CD3-CD8-GZMA-CD27 | 0.309 | 5 |
| High (top 20%) | B02_CD19-CD38-CD27-ICAM1 | M05_CD163-HLADR-CD16 | 0.298 | 5 |
| High (top 20%) | E03_EpCAM-LAMP1-B7H3 | E16_EpCAM-LAMP1-CD127-panRAS | 0.271 | 5 |
| High (top 20%) | F07_Fibronectin-IL1b | M06_CD68-LAMP1-HLADR | 0.250 | 5 |
| High (top 20%) | T01_CD3-GZMA-CD27 | M06_CD68-LAMP1-HLADR | 0.234 | 5 |
| High (top 20%) | M06_CD68-LAMP1-HLADR | M07_CD15-STING-PDL1 | 0.232 | 5 |
| High (top 20%) | B02_CD19-CD38-CD27-ICAM1 | T03_CD3-CD4-ICOS-CCR7 | 0.229 | 5 |
| High (top 20%) | B02_CD19-CD38-CD27-ICAM1 | B03_CD19_CD38_ICAM1 | 0.225 | 5 |
| High (top 20%) | M08_CD163-CD16 | M06_CD68-LAMP1-HLADR | 0.224 | 5 |
| High (top 20%) | T01_CD3-GZMA-CD27 | T02_CD3-CD8-GZMA-CD27 | 0.214 | 5 |
| Low (bottom 80%) | E04_EpCAM-STING-Her2 | E05_EpCAM-TCF7-Ki67-p53 | 0.295 | 5 |
| Low (bottom 80%) | E04_EpCAM-STING-Her2 | M04_CD68-IL1b-CD14 | 0.221 | 5 |
| Low (bottom 80%) | E10_EpCAM-CD11b-BetaCatenin-PDL1 | E05_EpCAM-TCF7-Ki67-p53 | 0.219 | 5 |
| Low (bottom 80%) | E04_EpCAM-STING-Her2 | B02_CD19-CD38-CD27-ICAM1 | 0.203 | 5 |
| Low (bottom 80%) | E16_EpCAM-LAMP1-CD127-panRAS | E14_EpCAM-BetaCatenin-panRAS | 0.198 | 5 |
| Low (bottom 80%) | M06_CD68-LAMP1-HLADR | T02_CD3-CD8-GZMA-CD27 | 0.190 | 5 |
| Low (bottom 80%) | N01_CD56 | NK01_CD56-CD38-HLADR | 0.189 | 5 |
| Low (bottom 80%) | T02_CD3-CD8-GZMA-CD27 | M06_CD68-LAMP1-HLADR | 0.189 | 5 |
| Low (bottom 80%) | B02_CD19-CD38-CD27-ICAM1 | T03_CD3-CD4-ICOS-CCR7 | 0.180 | 5 |
| Low (bottom 80%) | F04_Fibronectin-CCR7-Ki67 | F02_Fibronectin-SMA-ILb-CD15 | 0.177 | 5 |

**Table 4.b.** Spatial co-variation between B7-H3 and remaining panel markers across validation cohort cores. Mean Lee’s L value and standard deviation (SD Lees L) indicate the spatial co-variation between the B7-H3 anchor and the corresponding partner marker across tissue cores (n_cores). Proportion significant column denotes the proportion of the cores where the association was significant, and the robust column denotes whether significance was observed in more or less than 50% of cores. Positive Lee’s L values indicate positive spatial co-variation.

| Anchor | Partner | Mean Lees L | SD Lees L | n_cores | Proportion significant | Robust |
|---|---|---|---|---|---|---|
| B7-H3 | 4-1BB | -0.014 | 0.123 | 74 | 0.365 | <50% cores significant |
| B7-H3 | Bcl-2 | -0.052 | 0.117 | 74 | 0.216 | <50% cores significant |
| B7-H3 | Beta-catenin | -0.057 | 0.175 | 74 | 0.257 | <50% cores significant |
| B7-H3 | CCR7 | -0.023 | 0.120 | 74 | 0.338 | <50% cores significant |
| B7-H3 | CD11b | -0.019 | 0.142 | 74 | 0.257 | <50% cores significant |
| B7-H3 | CD11c | 0.081 | 0.126 | 74 | 0.676 | > 50% cores significant |
| B7-H3 | CD123 | -0.031 | 0.130 | 74 | 0.257 | <50% cores significant |
| B7-H3 | CD127 | -0.056 | 0.181 | 74 | 0.297 | <50% cores significant |
| B7-H3 | CD138 | -0.060 | 0.133 | 74 | 0.122 | <50% cores significant |
| B7-H3 | CD14 | 0.014 | 0.117 | 74 | 0.419 | <50% cores significant |
| B7-H3 | CD15 | -0.059 | 0.111 | 74 | 0.162 | <50% cores significant |
| B7-H3 | CD16 | 0.096 | 0.116 | 74 | 0.770 | > 50% cores significant |
| B7-H3 | CD163 | 0.016 | 0.089 | 74 | 0.473 | <50% cores significant |
| B7-H3 | CD19 | -0.023 | 0.134 | 74 | 0.284 | <50% cores significant |
| B7-H3 | CD20 | -0.060 | 0.108 | 74 | 0.176 | <50% cores significant |
| B7-H3 | CD27 | -0.049 | 0.126 | 74 | 0.243 | <50% cores significant |
| B7-H3 | CD3 | -0.015 | 0.122 | 74 | 0.338 | <50% cores significant |
| B7-H3 | CD31 | -0.004 | 0.091 | 74 | 0.432 | <50% cores significant |
| B7-H3 | CD34 | 0.001 | 0.118 | 74 | 0.500 | ≥ 50% cores significant |
| B7-H3 | CD38 | -0.042 | 0.112 | 74 | 0.203 | <50% cores significant |
| B7-H3 | CD39 | 0.165 | 0.125 | 74 | 0.865 | > 50% cores significant |
| B7-H3 | CD4 | 0.052 | 0.106 | 74 | 0.595 | > 50% cores significant |
| B7-H3 | CD40 | 0.069 | 0.165 | 74 | 0.581 | > 50% cores significant |
| B7-H3 | CD45 | 0.041 | 0.153 | 74 | 0.500 | ≥ 50% cores significant |
| B7-H3 | CD45RA | -0.093 | 0.177 | 74 | 0.162 | <50% cores significant |
| B7-H3 | CD56 | -0.022 | 0.091 | 74 | 0.270 | <50% cores significant |
| B7-H3 | CD68 | 0.007 | 0.135 | 74 | 0.419 | <50% cores significant |
| B7-H3 | CD8 | -0.017 | 0.060 | 74 | 0.216 | <50% cores significant |
| B7-H3 | CTLA4 | -0.040 | 0.122 | 74 | 0.189 | <50% cores significant |
| B7-H3 | EGFR | -0.046 | 0.192 | 74 | 0.270 | <50% cores significant |
| B7-H3 | EpCAM | -0.018 | 0.177 | 74 | 0.270 | <50% cores significant |
| B7-H3 | FABP4 | -0.086 | 0.116 | 74 | 0.162 | <50% cores significant |
| B7-H3 | FOXP3 | -0.057 | 0.147 | 74 | 0.162 | <50% cores significant |
| B7-H3 | Fibronectin | 0.041 | 0.143 | 74 | 0.527 | > 50% cores significant |
| B7-H3 | GITR | -0.018 | 0.123 | 74 | 0.324 | <50% cores significant |
| B7-H3 | GZMA | -0.003 | 0.108 | 74 | 0.351 | <50% cores significant |
| B7-H3 | GZMB | 0.006 | 0.119 | 74 | 0.446 | <50% cores significant |
| B7-H3 | HLA-DR | 0.076 | 0.112 | 74 | 0.662 | > 50% cores significant |
| B7-H3 | Her2 | -0.039 | 0.133 | 74 | 0.257 | <50% cores significant |
| B7-H3 | ICAM1 | 0.096 | 0.132 | 74 | 0.770 | > 50% cores significant |
| B7-H3 | ICOS | -0.018 | 0.116 | 74 | 0.351 | <50% cores significant |
| B7-H3 | IDO1 | -0.047 | 0.114 | 74 | 0.216 | <50% cores significant |
| B7-H3 | IL-18 | -0.045 | 0.123 | 74 | 0.257 | <50% cores significant |
| B7-H3 | IL-1b | -0.006 | 0.136 | 74 | 0.338 | <50% cores significant |
| B7-H3 | IgD | -0.002 | 0.118 | 74 | 0.351 | <50% cores significant |
| B7-H3 | Ki-67 | -0.015 | 0.104 | 74 | 0.257 | <50% cores significant |
| B7-H3 | LAG3 | -0.008 | 0.128 | 74 | 0.405 | <50% cores significant |
| B7-H3 | LAMP1 | -0.033 | 0.169 | 74 | 0.257 | <50% cores significant |
| B7-H3 | NF-kB p65 | -0.020 | 0.198 | 74 | 0.338 | <50% cores significant |
| B7-H3 | PD-1 | -0.030 | 0.155 | 74 | 0.324 | <50% cores significant |
| B7-H3 | PD-L1 | -0.061 | 0.176 | 74 | 0.257 | <50% cores significant |
| B7-H3 | PD-L2 | -0.039 | 0.138 | 74 | 0.230 | <50% cores significant |
| B7-H3 | SMA | 0.066 | 0.176 | 74 | 0.608 | > 50% cores significant |
| B7-H3 | STING | -0.072 | 0.093 | 74 | 0.122 | <50% cores significant |
| B7-H3 | TCF7 | -0.060 | 0.116 | 74 | 0.189 | <50% cores significant |
| B7-H3 | Tim-3 | 0.083 | 0.133 | 74 | 0.703 | > 50% cores significant |
| B7-H3 | VISTA | -0.019 | 0.117 | 74 | 0.338 | <50% cores significant |
| B7-H3 | Vimentin | 0.087 | 0.174 | 74 | 0.622 | > 50% cores significant |
| B7-H3 | iNOS | -0.041 | 0.131 | 74 | 0.243 | <50% cores significant |
| B7-H3 | p53 | -0.026 | 0.122 | 74 | 0.257 | <50% cores significant |
| B7-H3 | pan-RAS | -0.106 | 0.163 | 74 | 0.149 | <50% cores significant |

Lee’s L bivariate spatial association analysis showed B7-H3 exhibited consistent positive association with markers of immune exhaustion, myeloid activation, and stromal remodelling, including CD39, CD16, ICAM1 and vimentin across cores, independent of overall B7-H3 expression (Figure 6.c and Table 4.b; Supplemental Figure 5.b). This positions B7-H3 within potential exhausted T-cell and fibro-inflammatory stromal niches, consistent with the neighbourhood and proximity patterns described above.

#### B7-H3 expression associates with recurrent local cellular interactions

To determine whether the spatial relationships above extend to cellular interaction, we performed pseudo cell-cell interaction across B7-H3 ranked patient groups. Analysis revealed significant recurrent spatial cell-cell networks across B7-H3 expression states, including B7-H3^high^ tumours showing positive enrichment between fibroblasts and macrophages (Figure 6.d and Supplemental Figure 5.c). Aggregation confirmed a shift in edge composition, with stroma-immune interactions overrepresented among B7-H3^high^ edges, while epithelial-epithelial interactions were enriched among B7-H3^low^ edges (Figure 6.e). Interactions with near-zero differential enrichment indicated conserved baseline architecture across conditions, confirming a selective restructuring. Collectively, this hierarchical spatial analysis strategy supports a spatial model in which B7-H3 seems to function as key protein of the PDAC tumour environment rather than simply acting as a marker of individual cell states, consistently stratifying tumours into two reproducible states; B7-H3^low^ epithelial-centric and B7-H3^high^ fibro-inflammatory (Figure 6.f).

### Orthogonal RNA spatial profiling validates *CD276* (B7-H3) associated tumour patterns

To test whether the spatial architecture observed at the protein level reflects a transcriptionally relevant tumour state, we performed 6K RNA CosMx profiling on a serial section from the PDAC validation cohort (Figure 1.a-b). RNA-based cell typing was performed to recapitulate important cell types observed in the protein (Figure 7.a and Supplemental Figure 6.a). *CD276* (B7-H3) transcript enrichment mirrored protein expression, concentrating predominantly within malignant epithelium and fibroblast subtypes (Supplemental Figure 6.a-c). Univariate Ripley’s K analysis demonstrated non-random clustering of immune cells, inflammatory fibroblast and macrophage states within matched *B7-H3*^high^ cases, as well as clustering of activated epithelial populations within *B7-H3*^low^ cases (Figure 7.b). Lee’s L revealed consistent positive spatial dependency between *CD276* (B7-H3) and immune transcripts (*MRC1, GRAMD1B and IFI27*) and fibro-inflammatory transcripts (*MMP14, FLNA and BGN*) (Figure 7.c). These results recapitulate the fibro-immune/epithelial-intrinsic states identified in the protein dataset, as well as reproducing the spatial interaction partners, confirming that *CD276* (B7-H3) marks stable spatial states conserved across molecular layers.

**Figure 7:**
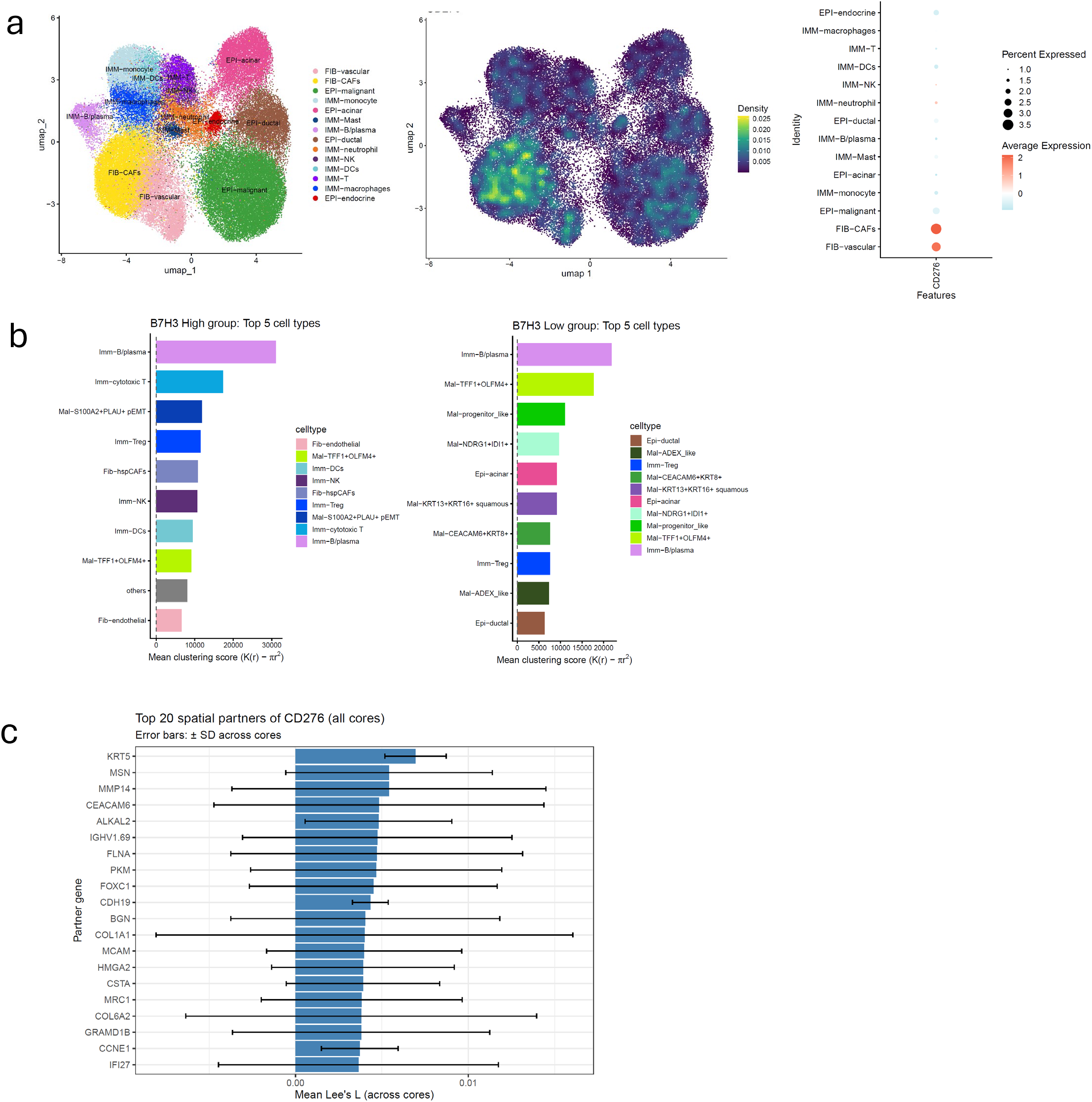
RNA panel reproduces CD276 (B7-H3) cell-type enrichment and spatial structure seen in protein. **a).** UMAP representing parent cell types generated from a serial section of the treatment naive validation PDAC cohort using single cell spatial transcriptome CosMx 6K panel (left), UMAP showing normalized *CD276* (B7-H3) expression, and a dot plot showing the average expression of *CD276* (B7-H3) across parent cell types, **b).** Ranked bar chart showing Univariate Ripley’s K mean degree of clustering of top final RNA subclusters split across *CD276* (B7-H3) ranked expression, coloured by cell type. Mean delta K degree of clustering = represents the difference between the observed spatial pattern and the theoretical random distribution. Observed K = the actual count, Exact CSR = complete spatial randomness, **c).** Ranked bar chart showing mean observed Lee’s L across cores (range = -0.1 to 0.2), with *CD276* (B7-H3) as the anchor and all other markers as the partner. Top 20 partners are shown.

## Discussion

Pancreatic cancer remains characterised by its dense fibrotic landscape, with the interactions between malignant cells, stromal cells and the immune population helping drive tumour progression and therapeutic resistance (5–7). Although B7-H3 has emerged as a promising immunotherapeutic target across multiple solid tumours, its biological role in the spatial organising of the pancreatic cancer tumour microenvironment remains poorly understood (19–21, 25, 34). To help resolve these complexities, we integrated regional spatial profiling with single cell spatial proteomics and transcriptomics across two independent treatment naïve resected patient cohorts. Our study demonstrates that B7-H3 is not simply overexpressed within individual cellular compartments but instead helps define reproducible tumour states. Across complementary analytical approaches and using orthogonal data across different molecular platforms, B7-H3^high^ tumours consistently present with fibro-inflammatory spatial organisation characterised by stromal and immune interactions. In contrast, B7-H3^low^ tumours retained a predominantly epithelial centred architecture with limited stromal/immune interactions. Together, these findings position B7-H3 as a multi-layered/multi-scale spatial organiser of PDAC, rather than solely an immune checkpoint molecule.

Regional spatial analysis suggests B7-H3 expression reflects distinct biological programmes, extending beyond tumour cell expression alone. Consistent with literature reports linking elevated B7-H3 expression to poor prognosis in PDAC (19), patients with elevated B7-H3 expression significantly correlated with poor disease specific survival within the protein and the transcriptomic landscape. Pathway analysis further reveals that patients within the top 20^th^ percentile of CD276 *(B7-H3)* expression were characterised by activation of fibroblast activation, extracellular remodelling pathways and inflammatory immune pathways, whereas the bottom 80^th^ percentile of patients retained epithelial associated signalling programmes with limited immune/stromal pathway activation. A similar phenomenon has also been seen in colorectal cancer tumours, with CD276 *(B7-H3)* enriched tumours significantly correlating with increased fibrotic and immune differentially expressed gene signatures (35). These findings suggest that CD276 *(B7-H3)* expression may represent a tumour microenvironment phenotype rather than a cellular biomarker.

Single-cell spatial protein profiling demonstrated that B7-H3 expression is elevated across multiple malignant epithelial, fibroblast and myeloid populations, and remains concentrated within highly organised multicellular neighbourhoods. Previous studies have reported multiple sites of localisation of B7-H3, with variable expression within tumour, stromal and immune populations, our findings provide a potential explanation for these discrepancies by demonstrating that cellular lineage alone does not adequately define B7-H3 biology (36–38). B7-H3 preferentially localises within conserved fibro-inflammatory neighbourhoods composed of interacting stromal and immune populations as seen in the results generated in figure 5. This spatial context is largely lost in conventional analysis approaches, highlighting the importance of preserving tissue architecture when investigating immune checkpoint biology.

Importantly, multiple independent analyses converged upon the same biological interpretation. Neighbourhood analysis identified recurrent fibro-immune enrichment with B7-H3 elevated micro-neighbourhoods, whereas epithelial cells were consistently depleted. Distance mapping further demonstrated that these micro-neighbourhoods were not isolated structures, gradient analysis showed fibro-inflammatory spatial domains preferentially enriched immediately adjacent to B7-H3 enriched micro-neighbourhoods, while epithelial cell presence progressively increased with distance. These observations suggest that B7-H3 associated function as local centres from which tissue composition transitions towards more epithelial-dominant architecture, supporting the concept of spatial microenvironment states rather than discrete cellular aggregates.

This microenvironment model was reinforced by complementary point pattern statistical analysis. Ripley’s K demonstrated preferential clustering of activated epithelial populations within B7-H3^low^ tumours, but clustering of inflammatory fibroblast, macrophages and innate immune populations within B7-H3^high^ tumours. Neatest neighbour analysis identified selective short-range coupling between activated fibroblast and myeloid subsets, whiles Lee’s L spatial co-variation consistently associated B7-H3 with markers of immune exhaustion, stromal activation and extracellular matrix remodelling. Although each analysis method interrogates distinct aspects of tissue organisation, they all independently identified the same fibro-inflammatory architecture surrounding B7-H3 expression, providing robust evidence that these observations reflect true biologically organised spatial programmes rather than stochastic variation. This type of spatial architecture has previously been detected in some extent in lung cancer brain metastases, demonstrating that the localisation and integration of B7-H3 far outweighs density metrics when associating with prognosis, indicating that these complex spatial structures are biologically important (39).

One of the strongest pieces of evidence for this model comes from the interaction network analysis of figure 6. It reveals selective rewiring of cell-cell interaction topology from stromal-immune, particularly myeloid-fibroblast interactions, and immune-immune interactions in B7-H3^high^ samples to epithelial-epithelial in B7-H3^low^ samples, rather than simply demonstrating differences in cell abundance. Importantly, this rewiring remained consistent across multiple spatial interaction radii tested (from 15-25μm), indicating that these results are robust to modelling parameters, representing a stable feature in PDAC architecture, rather than artefacts of network construction.

The reproducibility of these findings across different spatial platforms further validates the described B7-H3 reorganisation model. Transcriptome profiling using the CosMx 6K panel recapitulated the dominant biological observations identified using single cell spatial proteomics. This convergence between spatial proteomic and transcriptomics strengthens the confidence that the described phenomenon represents genuine biological features of PDAC rather than platform specific or stochastic observations.

From a translational perceptive, these findings have important implications for the development of B7-H3 targeted therapies. There are currently multiple therapeutic strategies targeting B7-H3 being trialled across multiple solid tumours including antibody-drug conjugates (22, 23). Our data suggest that therapeutic response may depend not only on B7-H3 expression, but also on the spatial microenvironment in which B7-H3 resides. Patients with B7-H3 enriched, fibro-inflammatory tumour microenvironment may represent a biologically distinct subgroup with the highest likelihood of benefiting from B7-H3 directed approaches. Conversely, patients with B7-H3 low expression may benefit from alternative or combination therapeutic strategies. It is important to note that B7-H3 inhibition would likely only be applicable to patients with previously treated, unresectable cancer (22, 23). The success of KRAS inhibitors in the PDAC landscape may offer a potential avenue for combined immunotherapy with B7-H3 inhibitors.

This study has several limitations that should be acknowledged when interpreting the findings. First, all spatial analysis presented here are observational and therefore cannot establish causal interactions between B7-H3 and neighbouring cellular populations, functional perturbation experiments would be required to fully determine whether B7-H3 actively orchestrates microenvironment states or reflects a broader tissue remodelling program. Second, spatial proximity and co-occurrence do not necessarily imply direct molecular communication, establishing receptor-ligand pairings would be required, incorporating assays including proximity labelling strategies, something that has already been demonstrated in B7-H3 (40). Third, the panel used for the single-cell spatial protein experiment has limited coverage of epithelial lineage markers, which restricts the resolution of the epithelial cellular states named. Fourth, this study has been performed using TMA cores to increase the number of patients. To fully establish the B7-H3 associated fibro-inflammatory state within the spatial context, further work would be required in a bespoke whole section validation cohort. Finally, current clinical trials are focused of previously treated, advanced, unresectable PDAC. Further work would need to be performed, ideally, using a longitudinal cohort encompassing sections from biopsy, treated primary and treated metastatic patients to validate B7-H3 associated biology.

In conclusion, this study establishes B7-H3 as a multiscale organiser of the pancreatic tumour landscape rather than simply an immune checkpoint molecule. By integrating regional spatial profiling with single cell spatial proteomics and transcriptomics, we demonstrate that B7-H3 consistently defines fibro-inflammatory states characterised by coordinated epithelial, stromal and immune interactions. These programmes were reproducible across analytical methodology and molecular modalities, supporting their biological robustness. This work provides a spatial systems framework for understanding immune checkpoint biology embedded within tissue architecture. As spatial technologies become increasingly integrated into translational research, understanding the complex, spatially involved, environment in which biomarkers such as B7-H3 may improve patient stratification, guide therapeutic strategies, and ultimately inform more effective treatment strategies for pancreatic cancer, something which is desperately needed.

## Methods

### Ethics

Approval for these samples was obtained by the West of Scotland Research Ethics Committee and the NHS QEUH Biorepository. All clinical data is covered under Caldicott approval.

### Patient Cohort

A discovery treatment naïve PDAC tissue microarray (TMA) cohort (n=28) who underwent surgical resection between 2010 and 2017 was profiled using immunohistochemistry and regional spatial proteomics. Each case was represented by approximately 4×0.6 mm cores. Histologically, this cohort was predominantly epithelial-rich.

An independent treatment naïve PDAC TMA (n=79) who underwent resection between 1992 and 2011 was used as a validation cohort, and profiled using regional spatial transcriptomics, single cell spatial protein and single cell spatial transcriptomics. Each case was represented by a 1×0.6 mm core. This cohort displayed mixed histological architecture, including both tumour-dominant and stroma-rich cores.

All samples were formalin-fixed paraffin-embedded (FFPE) tissues with linked clinicopathological annotation (Supplemental Methods Table 1-2). All samples underwent surgical resection in the Glasgow Royal Infirmary (GRI).

### Regional Spatial Technologies

Regional spatial transcriptomic and proteomic profiling was performed using the GeoMx® Digital Spatial Profiler (DSP) platform. Assays were performed according to the manufacturer’s protocols, with minor modifications (Supplemental Methods 3). Region of interest (ROI) selection was performed blinded to clinical outcome.

#### Immune-Oncology Protein Panel assay

The GeoMx® Immuno-Oncology (IO) protein panel (60 targets) was applied to the discovery cohort as part of a NanoString technology access programme. The morphology panel staining included Syto13, Pan-cytokeratin (PanCk), CD3, and αSMA. A total of 48 geometric ROIs were profiled across 27 patients each with a diameter of 660μm. Within each ROI, compartment-specific Areas of illumination (AOI) were defined using the GeoMx DSP Control Centre native pixel-threshold segmentation algorithm: epithelial AOIs were defined by PanCk+ve staining (n=48), and tumour microenvironment (TME) AOIs were defined by PanCk-ve regions (n=48).

#### Human Whole Transcriptome Atlas (WTA) Panel assay

The GeoMx® Human Whole Transcriptome Atlas (WTA), (18,677 genes), was applied to the validation cohort. The morphology panel comprised Syto13 and PanCk (GeoMx morphology kit; GMX-RNA-MORPHHST-12), supplemented with in-house oligo-conjugated antibodies against CD45 (M087629-2, Thermo Fisher Scientific) and αSMA (53-9760-82, Thermo Fisher Scientific) (Supplemental methods 3). ROI selection was performed using the GeoMx DSP Control Centre native pixel-threshold segmentation algorithm. A total of 63 geometric ROIs were profiled at 660μm. Within each ROI, compartment-specific AOIs were defined as: epithelial (PanCk+; n=56), fibroblast-enriched (αSMA+; n=52), immune-enriched (PanCk-αSMA-; n=56), and mixed core (n=3). Multiple AOIs could be segmented per ROI where tissue architecture permitted.

### Single Cell Spatial Technologies

CosMx™ Spatial Molecular Imaging (SMI) was performed on serial sections from the validation cohort using the Immune-Oncology protein panel and the 6K Human Discovery RNA panel. The pilot neoadjuvant cohort was profiled using 6K Human Discovery RNA panel. Assays were performed according to the manufacturer’s protocol with tissue-specific optimisation of antigen retrieval and Proteinase K digestion where required (Supplemental methods 4). Fields of view (FOVs) were placed using a grid-based tiling strategy to cover entire cores or tissue regions. The validation cohort acquired 303 FOVs (n=74) using the IO protein panel, and 134 FOVs were acquired (n=48) using the 6K panel. 1,489 FOVS were acquired for the pilot neoadjuvant cohort (n=4).

### Spatial Data pre-processing

All data preprocessing was performed as described in supplemental methods 6a-e

### Statistical analysis

All P-values are reported to 3 decimal places and were adjusted for multiple testing using the Benjamini-Hochberg false discovery rate and indicated by FDR. Significance was set to 0.05 unless otherwise indicated.

### Kaplan Meir analysis

Log-rank survival and univariate cox regression analysis was performed throughout to establish associations for disease specific survival (DSS) and recurrence free survival (RFS) with RStudio using survminer and survival packages. Multiple hypothesis testing was applied.

### Regional spatial statistical analysis

#### Spearman correlation

Concordance between IHC and matched regional protein markers CD3, CD8 and CD68 was performed with Spearman Rank correlation using the ggplot2 R package.

#### Differential expression analysis (DEA)

Differential expression analysis (DEA) was carried out by EdgeR and EnhancedVolcano R package. Statistical significance was set to log2 fold change ±1.5 along with FDR.

#### Paired Violin Boxplots

Regional B7-H3 protein expression was compared between paired AOIs using a linear mixed-effects model to account for repeated sampling within patients. Expression values were log10-transformed to reduce skew. Segment was modelled as a fixed effect, and patient identity was included as a random intercept to account for within-patient correlation arising from multiple cores per patient. Statistical significance of the segment effect was assessed using an F-test derived from the mixed model. Data visualization was performed using violin and box plots with overlaid individual core measurements and paired patient means. All analyses were conducted in R using packages lme4 and lmerTest, and FDR was reported.

#### Correlation heatmap

Spearman correlations along with FDR were computed between B7-H3 and all protein markers across TME compartments.

#### Principle Component Analysis (PCA)

PCA was performed on normalized data using R package prcomp. PC1 and PC2 were used to visualise global transcriptomic variance across AOIs, points were coloured by segment identity.

#### Gene Set Enrichment Analysis (GSEA)

Gene set enrichment analysis (GSEA) was generated by the fgsea R package, using the GeoMx filtered pathways. Statistical significance was set to normalised enrichment score (NES) 1.5/-1.5 and FDR reported. Visualization was performed using ggplot2, with top 10 significant pathways per group presented.

#### Spatial Deconvolution (SpatialDecon)

Immune cell deconvolution was performed using estimates the immune cells present within the bulk transcriptomic context using validated single cell signatures and the SpatialDecon R package. Immune cell count per 100 cells was generated, and B7-H3 ranked expression groups were compared. Statistical analysis was performed using Wilcoxon test.

### Single cell protein spatial statistical analysis

#### Protein Cell typing

Cell identities were defined using an iterative graph-based clustering strategy implemented in the Seurat R package. Following normalisation and PCA, a nearest-neighbour graph was constructed and Louvain clustering performed. Clustering resolution was progressively increased on the full dataset (without sub-setting) to refine cellular organisation from broad lineages to stable parent cell types. Cell types were first defined using differential protein expression assessment using FindAllMarkers (MAST test; minimum fraction of cells = 25%, FDR <0.05). Signatures generated were investigated for canonical markers associated with key cell types, then images were reviewed for spatial localisation, and concordance with matched H&E and multiplex immunofluorescence images. Clusters were reviewed by a pathologist where appropriate

To achieve higher-resolution subtype definition, selected parent populations (epithelial cells, fibroblasts, macrophages, T cells, B cells,) were extracted and re-clustered independently using the same framework at varying resolutions. If the signature generated was clearly mixed cell types, this was visually interrogated and manually re-named if appropriate. A selection of the top enriched markers were incorporated into the final subtype naming conventions (e.g., E01_EpCAM-marker1-marker2).

At lower resolution, cells split into major lineages defined by protein expression profiles: epithelial, lymphoid, myeloid, stromal, and a stromal-vascular mixed compartment. Further resolution identified coherent parent populations within these lineages, including epithelial tumour and endocrine cells; T cells, B cells, and NK cells within the lymphoid compartment; macrophages, monocytes, neutrophils, and tumour-associated macrophages (TAMs) within the myeloid lineage; and fibroblasts within the stromal compartment. A stromal-vascular mixed population was retained as two mixed groups due to panel limitations and overlapping CD31 signal within fibroblast-rich regions, preventing robust endothelial-fibroblast separation, as demonstrated in Supplemental Figure 4.a and Supplemental Figure 4.d. Additional to filtering above, red blood cell and endocrine cell clusters were removed for downstream analysis, resulting in a total of 167,740 cells.

#### Smoothened and ranked B7-H3 cell types

To identify B7-H3 enriched clusters, spatially smoothed expression UMAPs were generated using the Hieratype R package. Smoothed intensity distributions were used to visualise local enrichment patterns across fine-grained subclusters and confirm spatial coherence of signal. Subclusters were then quantitatively ranked by mean B7-H3 protein expression. A subcluster was defined as B7-H3 enriched if its mean expression ranked among the highest across all subclusters and ≥70% of cells within that subcluster exhibited elevated normalized B7-H3 expression, ≥50, relative to the global cellular distribution.

#### Macro niche generation

To capture tissue-scale architectural states, we generated macro niches based on local cell-type composition within a fixed physical radius. For each cell, we identified neighbouring cells within a 50 µm radius using a fast k-nearest neighbour search (RANN), querying up to 500 neighbours per cell and retaining only those within the radius threshold.

A neighbourhood composition vector was computed for each focal cell, counting the number of neighbouring cells assigned to each parent lineage, producing a per-cell composition matrix. Composition vectors were centred and scaled, denoised (top 10 principal components), and clustered using k-means (k = 10 and fixed random seed) to define macro-scale structural niches.

#### Micro neighbourhood generation and analysis

To generate micro niches, we performed graph-based clustering in protein expression space using Seurat. A shared nearest neighbour (SNN) graph was constructed from Harmony embeddings (first 30 components, k = 10 nearest neighbours, Euclidean distance and Jaccard edge pruning = 0.067). The resulting SNN graph was used for clustered using Leiden community detection (resolution = 1.0), yielding expression-defined micro-neighbourhood clusters. For each patient core, mean B7-H3 expression was calculated within each micro-neighbourhood. Micro neighbourhood enrichment was defined as the deviation of a patient mean B7-H3 from the overall mean across all patient neighbourhood combinations, and overall differences in B7-H3 across micro neighbourhoods were assessed using a Kruskal-Wallis test.

Supervised comparison of B7-H3 associated micro neighbourhoods were stratified into ‘B7-H3 high’ and ‘B7-H3 low’ states using effect-size thresholds (top 20% and bottom 20%), with minimum cell-count filter (n= 200). Differences in composition between B7-H3 high and B7-H3 low neighbourhood were tested using Wilcoxon rank-sum tests.

#### Spatial Domain generation

Spatial domains were computed to quantify how tumour microenvironment composition changes with distance from B7-H3 high micro-neighbourhoods. For each patient, the Euclidean distance from each cell to its nearest B7-H3 high micro-neighbourhood cell was calculated. Distances were binned into 5 µm and patient-centred cell-type proportions were summarised per distance bin. Results were reported for five main microenvironment compartments made up of; epithelial-rich (epithelial parent cell type), fibro-inflammatory (fibroblasts, Macrophage and TAMs parent cell types), immune-rich (T cell, B cell, monocyte, neutrophil NK parent cell types), and mixed regions (all other cell types).

#### Point Pattern: Univariate Ripley’s K and Bivariate Neighbourhood

Spatial point-pattern analysis was performed using univariate Ripley’s K and bivariate nearest-neighbour statistics implemented in R using the SpatialTIME package. Analyses were conducted per tissue core then averaged. For univariate analysis, Ripley’s K was calculated for each cell phenotype across increasing spatial radii (r = 0–0.3 mm, step 0.01 mm, 199 permutations, translational edge correction). Observed K values were compared against a null model of complete spatial randomness (CSR). The degree of clustering at each radius was defined as:

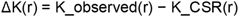

where positive values indicate clustering and negative values indicate dispersion relative to random expectation. ΔK(r) was averaged across 10–50 µm, yielding a mean per phenotype per core.

For bivariate analysis, nearest-neighbour distributions between B7-H3 expressing phenotypes and other phenotypes were computed (r = 0–0.6 mm, step 0.01 mm, 199 permutations, ‘rs’ edge correction). CSR distributions were generated via permutation of phenotype identities while preserving spatial coordinates. Spatial association was quantified as:

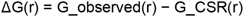

where positive values indicate enriched proximity and negative values indicate spatial avoidance. Mean ΔG values were averaged across 10–50 µm radius window to summarise phenotype-specific spatial coupling.

#### Spatial autocorrelation and spatial co-variation statistics – Moran’s I and Lee’s L

To quantify spatial autocorrelation of marker expression within cores, Moran’s I was computed for each protein marker using a core-specific neighbour graph. Moran’s I and associated p-values were computed per marker per core and were adjusted using FDR. Summary statistics were generated by aggregating Moran’s I across cores per marker.

To quantify co-variation between B7-H3 and other protein markers, we computed Lee’s L per tissue core using the spots R package. Statistical significance was assessed using a permutation framework (n = 500 permutations per core) in which B7-H3 expression values were randomly permuted across cells while preserving spatial coordinates and the neighbourhood graph. P-values were calculated as the proportion of permuted Lee’s L values greater than or equal to the observed statistic for each partner marker. Results were summarised across cores as mean Lee’s L with standard deviation and the proportion of cores passing FDR for each partner marker.

#### Spatial interaction network modelling

Cell-cell spatial interaction networks were constructed per B7-H3 ranked groups. For each patient core, an undirected proximity graph was built (r=20µm), and edges were restricted to within-patient links, with multi-edges or self-loops removed. Observed cell-cell edge counts were computed per patient, and expected counts were derived from within-patient phenotype frequencies under a random mixing null model proportional to cell-type abundance. Expected edge frequencies were aggregated after within-patient normalization. Enrichment was defined as observed/expected edge frequency.

Statistical support for non-random co-occurrence was assessed by comparing observed counts to the composition-matched null model using Fisher-style contingency statistics. Spatial interaction networks were inferred separately in B7-H3 ranks using pairwise co-occurrence statistics for all phenotype pairs. We counted how often each phenotype appeared in the significant enriched edge set and retained phenotypes participating in ≥3 significant edges. The top phenotypes per group were selected and combined to form a fixed node set, enabling direct comparison between B7-H3 ranked groups.

To quantify networks, significant enriched edges were extracted independently in B7-H3 ranks. Edges were merged across ranks to classify interactions as high-only, low-only, or shared. For shared edges, differential strength was defined as:

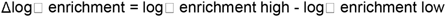

Positive values indicate stronger enrichment in B7-H3^high^ tumours.

To visualise analysis, a differential network was constructed from top-enriched edges using R packages igraph and ggraph.

### Single cell transcriptomic spatial statistical analysis

#### Cell typing and Path annotations – Validation naïve PDAC cohort

Cell-type annotation was performed using the CosMx 6K transcriptomic discovery panel using an iterative strategy integrating unsupervised transcriptomic clustering, marker-based annotation, spatial mapping and pathologist-guided histological review. After scVI integration, Leiden unsupervised clustering was performed on the scVI latent representation at a resolution of 1.0. Cluster-enriched marker genes were then used to generate an initial broad annotation of major cell classes. These preliminary annotations were visualised to assess their tissue distribution. In parallel, post-CosMx H&E images were reviewed by a pathologist, with manual annotation focused on vessels, nerves, normal epithelium and malignant epithelium. H&E-based annotations were transferred onto the corresponding CosMx images. The resulting cell IDs were then used to update the initial transcriptome-based annotations. Finally, iterative hierarchical subtype annotation was performed within the major cell classes to generate the final cell-type labels. The final annotations were spatially mapped and reviewed by a pathologist.

## Supporting information

Supplemental Results Figures

## Data and code availability

Data is available on request to the corresponding author

## Acknowledgements

We extend our gratitude to the patients and their families. We acknowledge the Academic unit of surgery at the Glasgow Royal Infirmary. We also acknowledge the Translation Tissue Service Glasgow, the Cancer Research UK Scotland Institute Histology Department, and the NHS Greater Glasgow and Clyde biorepository for helping with tissue preparation for this study.

## Funding

Assya Legrini and Nigel Jamieson were supported by Cancer Research UK (C55370/A25813), Pancreatic Cancer UK Early Detection grant (EDG2023-24_Jamieson) and Pancreatic Cancer UK Research Innovation Fund (RIF2023-24_Jamieson). Joanne Edwards was supported by Medical Research Council (MR/R502327/1), Greater Glasgow and Clyde endowment (306620–01), Cancer Research UK Scotland Centre (CTRQQR-2021\100006), the Beatson Cancer Charity (24-25-045), Beatson Cancer Charity (330619–01) and Chief Scientific Office (TCS/22/02). Kathryn Pennel was supported by Chief Scientific Office (EPD/22/13).

## Competing interests

None

## Supplemental information

Document S1 – supplemental methods text

Document S2 – supplemental results figure

Document S3 – supplemental tables

Document S4 – supplemental table 1

Document S5 – supplemental table 2

## Notes

### Competing Interest Statement

The authors have declared no competing interest.

