## Supplemental Methods Text for "B7-H3 Orchestrates Fibro-Immune Spatial states in Pancreatic Cancer"

### Supplemental Methods 1: Immunohistochemistry

The pilot cohort was stained on the Dako PT Link pre-treatment module and Autostainer Link 48 platform (Agilent Technologies) for CD3, CD8, CD68, and CD163 (Supplemental Methods Table 4). Antigen retrieval conditions were performed according to manufacturer-recommended protocols (Supplemental Methods Table 4). Following primary antibody incubation, detection was carried out using a horseradish peroxidase (HRP)-based polymer detection system with 3,3′-diaminobenzidine (DAB) chromogen. Slides were counterstained with haematoxylin, rinsed, dehydrated through graded ethanol, cleared in xylene, and mounted using DPX mountant (SEA-1300-00A, CellPath).

The section was scanned at 20X magnification using the Leica Aperio AT2 slide scanner (Leica Biosystems). Digital image analysis was performed using HALO® (Indica Labs, Albuquerque, USA). All cores were manually reviewed for quality prior to analysis; cores exhibiting blurring, tissue loss, or inadequate staining were excluded.

Marker-specific intensity thresholds were defined within HALO to classify staining as weak, moderate, or strong at the cellular level. For each core, a weighted histoscore (H-score) was calculated as:

H-score = (1 × % weak) + (2 × % moderate) + (3 × % strong)

yielding a continuous score ranging from 0 to 300. Mean H-scores were calculated per patient across available cores.

### Supplemental Methods 2: Regional omics assays

Prior to staining, FFPE sections were re-baked overnight at 60°C. Antigen retrieval was performed using the Leica BOND™ system (Leica Biosystems, Wetzlar, Germany) with ER2 buffer (HIER, 20 minutes). For RNA assays, enzymatic digestion was performed with recombinant Proteinase K (AM2546, Invitrogen) at 1 μg/mL for 15 minutes at 37°C.

For transcriptomic assays, samples underwent overnight in situ hybridisation (ISH) at 37°C using oligonucleotide probe panels. For protein assays, oligo-conjugated antibody panels were applied according to manufacturer specifications.

A morphology-based mini multiplex immunofluorescence (mIF) panel was used to visualise tissue architecture and guide ROI selection. Antibody–fluorophore conjugation was performed using Alexa Fluor™ labelling kits according to the manufacturer’s instructions. Conjugated antibodies were validated and approved for use by NanoString prior to assay deployment. CD45 and αSMA were applied at 1:200 dilution for 1 hour at room temperature, and Syto18 and PanCk were applied as recommended. Regions of interest (ROIs) were defined in the GeoMx DSP Control Center software, and areas of illumination (AOIs) were segmented based on marker-defined compartments using native thresholding algorithms. Oligonucleotide tags were collected and quantified using the nCounter® platform (NanoString) for protein assays, and sequenced on the Illumina NextSeq 1000/2000 platform for RNA assays.

### Supplemental Methods 3: Single cell omics assay

Sections were mounted within the gasket area (approximately 20 mm × 15 mm) on Leica BOND PLUS slides and baked overnight at 60°C. Pre-treatment was performed on the Leica BOND™ platform. Heat-induced epitope retrieval was carried out using ER1 buffer for 20 minutes at 100°C.

For RNA assays, tissue permeabilization was performed using recombinant Proteinase K (AM2546, Invitrogen) at 3 μg/ml for 15 minutes at 37°C. Incubation frames (CMX-FCA, NanoString) and fiducial markers (CMX-FFPE-SP-P, NanoString) were applied to enable spatial registration. Post-fixation and Sulfo-NHS-Acetate treatment (26777, Thermo Fisher Scientific) preceded overnight in situ hybridisation at 37°C. Stringent washes were performed according to manufacturer specifications.

For protein assays, a ready-to-use CosMx Immune-Oncology panel was applied. Nuclear staining was performed using DAPI (1:40 dilution). Cell segmentation markers CD298/B2M (1:50 dilution) were applied together with lineage markers including PanCk, CD45, and CD3 (1:25 dilution). For RNA assays (6K and WTA), nuclear staining (DAPI, 1:40) and segmentation markers CD298/B2M (1:50), PanCk, CD45, and CD68 (1:25) were applied.

Slides were processed on the CosMx™ instrument using manufacturer-recommended acquisition settings. Pre-bleaching configuration C and cell segmentation configuration A (human tissue profile) were used. Cell segmentation was performed using the CosMx modified Cellpose algorithm, integrating nuclear detection with membrane-guided cell expansion.

### Supplemental Methods 4: Spatial Data pre-processing

#### Supplemental 4.a: Regional Spatial Protein – Naïve Discovery PDAC cohort

Quality control (QC), normalisation, and target filtering of GeoMx® protein data were performed using the GeoMxTools R package. Housekeeping normalisation was applied using the geometric mean of GAPDH, Histone H3, and RPS6, as recommended for the Immuno-Oncology panel. Technical QC metrics were assessed according to NanoString manufacturer guidelines and included AOI surface area, nuclei count, binding density, negative control probe performance, and positive control linearity. All 98 AOIs passed predefined technical QC thresholds and were retained for analysis.

Targets were evaluated for signal-to-noise performance relative to matched negative control probes. Markers with signal-to-noise ratio (SNR) < 1.3 were flagged and reviewed; low-performing targets were removed where consistent low signal was observed across AOIs. Two TME AOIs were excluded from downstream analysis, resulting in 94 AOIs (tumour = 48 AOIs, TME = 46 AOIs).

#### Supplemental 4.b: Regional Spatial Transcriptomic – Naïve Validation PDAC cohort

Transcriptomic QC was performed using GeoMxTools following manufacturer-recommended thresholds, adjustments were made where indicated. Technical background was assessed using the geometric mean of negative probe counts per AOI. Given the elevated stromal content and FFPE-associated background typical of PDAC tissue, a negative probe geometric mean threshold of ≥4 counts was applied. AOIs falling below this threshold were flagged for inspection.

Minimum nuclei count threshold of 100 nuclei per AOI was applied to ensure adequate cellular representation. AOIs failing nuclei count or background QC were reviewed and excluded where appropriate. Full cores (n=3) were also excluded from downstream analysis. After QC filtering, 148 AOIs were retained for downstream analysis. Gene-level filtering was applied to remove low-abundance targets. Genes with expression below the median positive control probe count in >95% of AOIs were excluded. Due to imaging hardware constraints, focal tissue loss during processing and QC filtering, only 58/79 patients met inclusion criteria for final WTA analysis. Data normalisation was performed using the upper quartile (Q3) method, as implemented in GeoMxTools, to account for library size and technical variability across AOIs.

#### Supplemental 4.c: Single Cell Spatial Technologies

Raw CosMx™ data files, including Seurat objects and associated image files, were exported from AtoMx (version 2.3) for downstream analysis in RStudio. Morphology images were reviewed using Napari (v0.4.17) to assess overall tissue integrity, segmentation quality, and imaging artefacts prior to quantitative analysis.

##### Supplemental 4.d: Single Cell Spatial Protein Panel – Naïve Validation PDAC cohort

FOV images were manually reviewed for technical quality, including segmentation accuracy and staining integrity. Cores exhibiting tissue lifting, extensive blurring, segmentation failure, or poor staining, and cells that were identified as red blood cells were excluded prior to downstream processing.

Single-cell level quality control was subsequently applied to remove segmentation artefacts and low-quality cellular events. Cells were excluded if ≥50% of measured protein markers fell within the global 90th percentile of intensity (indicative of signal saturation or segmentation artefact), or if fewer than three markers were detected above the global 50th percentile, consistent with low-signal or debris-associated events. Cells with aberrant negative control probe values (<2 or >50) were removed to exclude background artefacts and non-specific signal. Segmented objects with an area >500 µm² were excluded to minimise inclusion of merged or improperly segmented cells. Following filtering, 296/303 FOVs and 170,923 cells (92% retention) were kept for downstream analysis.

Protein expression values were normalised using total-intensity scaling per cell to account for differences in overall signal capture. Normalised counts were subsequently arcsinh-transformed (cofactor = 5) to stabilise variance and compress high-intensity outliers, consistent with established cytometry-style workflows for multiplexed protein imaging. The transformed expression matrix was centred and scaled prior to dimensional reduction and clustering in Seurat.

##### Supplemental 4.e: Single Cell Spatial 6K RNA Panel – Naïve Validation PDAC cohort

Morphology-stained images were reviewed in Napari (v0.4.17) to assess tissue integrity, focus quality, and segmentation performance. Segmentation review resulted in approximately 5% of cells were flagged for potential segmentation artefacts, indicating high overall segmentation fidelity.

Single-cell quality control filtering was performed across all samples. Cells were excluded if total transcript counts were <20, segmented cell area exceeded 300 µm², or if the proportion of negative control probe counts exceeded 0.1, indicative of elevated background signal. After filtering, 100 FOVs were retained, comprising 73,568 cells (59.59% retention), with a mean transcript count per cell of 150.39.

Data normalisation and downstream analysis were performed in Seurat (v5.0.0). Counts were log-normalised using the default Seurat normalisation method (scale factor = 10,000). Highly variable genes were identified using the variance stabilising transformation (VST) method (n = 2,000 features), followed by centring and scaling of the expression matrix.

Data integration was performed using scVI implemented in scvi-tools (v1.4.0). Raw count data stored in the counts layer were used as input, and the AnnData object was registered using scvi.model.SCVI.setup_anndata with patient_label specified as the batch key. A random seed of 12345 was set for reproducibility. The scVI model was trained for 109 epochs, with other model parameters left as default. The learned scVI latent representation was used for nearest-neighbour graph construction, followed by Leiden clustering and UMAP visualisation.

Dimensionality selection was guided by inspection of the principal component elbow plot, which demonstrated an inflection point at approximately 25 components. To ensure robust representation of biological variance while avoiding overfitting to noise, the first 30 Harmony-corrected dimensions were retained for nearest-neighbour graph construction, clustering, and UMAP visualisation. Clustering stability across a range of resolutions was assessed, and the selected resolution (1.0) provided coherent, biologically interpretable clusters without excessive fragmentation.

### Supplemental Methods 5: B7-H3 ranked groups

Survival cut points were initially generated using the maximally selected rank statistics, which selects the statistically optimal threshold for splitting continuous data into high/low risk groups. This produced a 26%high/74%low split for stromal compartments (Table 1.a). However, we were specifically interested in the biological difference found within the most extreme B7-H3 expressors. To ensure that the percentile cut point chosen was not arbitrarily selected to maximise statistical significance, a sensitivity analysis was performed testing all percentile thresholds from the 50th to the 95th percentile in 5-percentile increments (Supplemental Methods Table 5). At each threshold, a log-rank test was performed comparing survival between the resulting High/Low groups and was adjusted using Benjamin-Hochberg. Cut points yielding fewer than 3 events in either group were excluded from testing.

Statistically significant survival separation (p-adjusted < 0.05) was observed across the multiple of tested cut points (50th, 55th, 75th, 80th percentiles), demonstrating that the prognostic signal of B7-H3 is robust and not dependent on a single threshold (Supplemental Methods Table 5).

Cut points at the 50th and 55th percentiles produced lower p values (p-adjusted = 0.021 and p-adjusted = 0.021, respectively). However, these splits classify 44-50% of the cohort as ‘High expressers’, conflating moderate and high expression and making the grouping biologically uninterpretable as an overexpressed phenotype (Supplemental Methods Table 5). The 75^th^ cut point also produced a lower p value (p-adjusted = 0.022); however, we were interested in exploring only the most extreme, significant B7-H3 expressors. The 80th percentile cut point preserves a clinically meaningful distinction between a minority overexpressed population and the remainder of the cohort. The final cut-off method was replicated for each experiment.
