## Supplemental Results Figures for "B7-H3 Orchestrates Fibro-Immune Spatial states in Pancreatic Cancer"

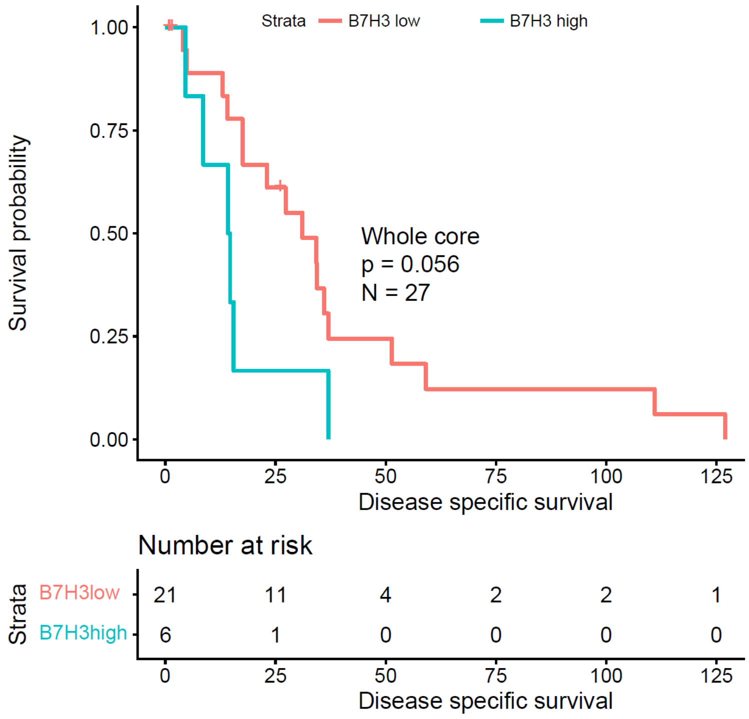


***Supplemental figure 1:*** *Regional immune landscape defines survival groups in PDAC*. Kaplan Meier curve (Log-Rank test) stratified by disease specific survival and associated with mean combined whole core B7-H3 expression.

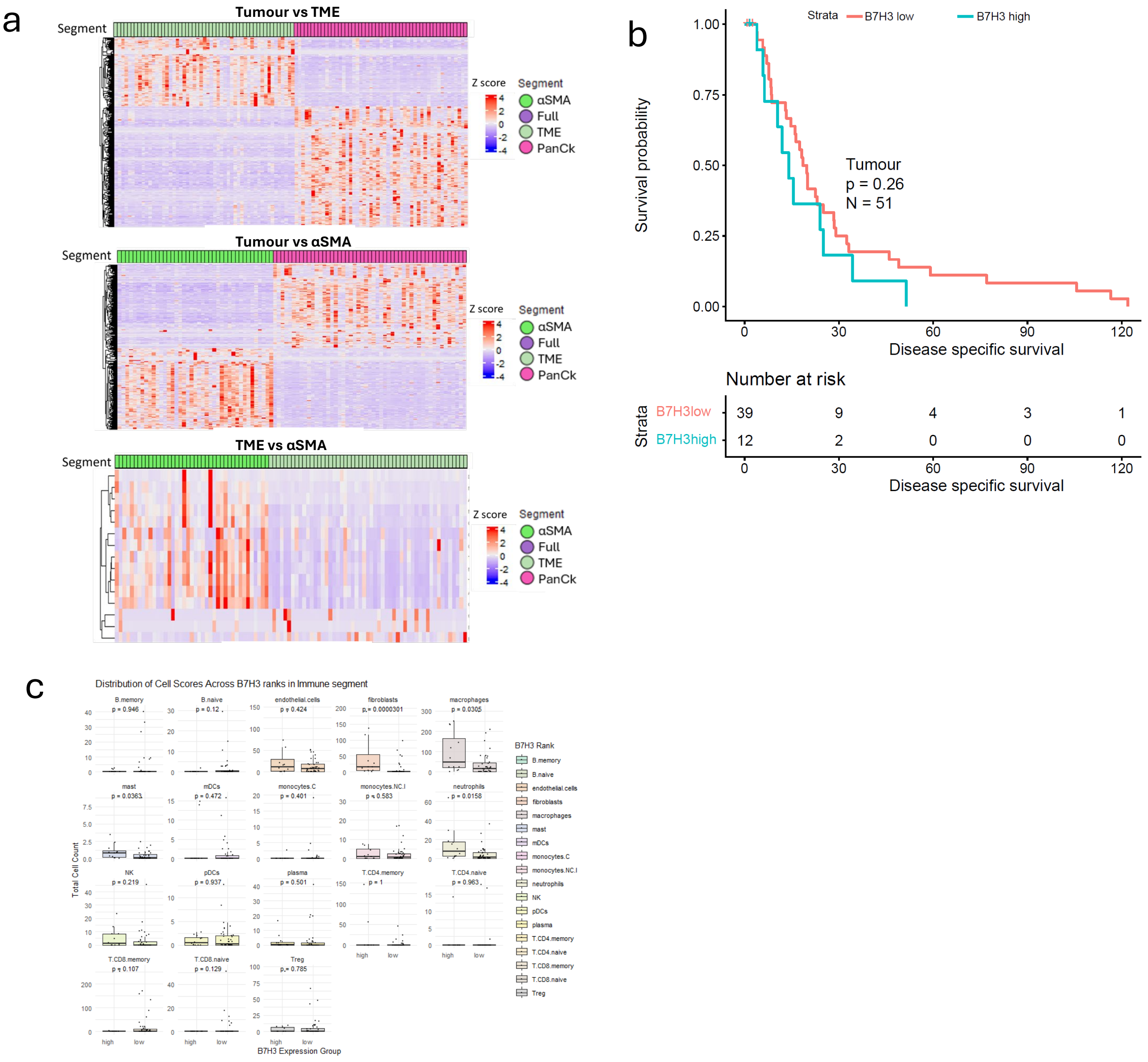


***Supplemental figure 2:*** *Regional protein profiling defines B7-H3 high as fibro-inflammatory stromal rich with immune suppression, and B7-H3 low maintains epithelial intrinsic states*

**a).** Differential expression heatmap showing significant differentially expressed genes in all segment comparisons. Comparisons carried out between Tumour vs TME, Tumour vs αSMA and TME vs αSMA, colours indicate the segments being compared, **b).** Kaplan Meier curves (Log-Rank test) stratified by disease specific survival and in tumour compartments of B7-H3 ranked patients, **c).** Boxplot showing Spatial immune cell deconvolution estimates of the proportion of immune cell within B7-H3 ranked patients in TME segments. Coloured by spatial deconvoluted cell type, p values generated by Wilcoxon test.

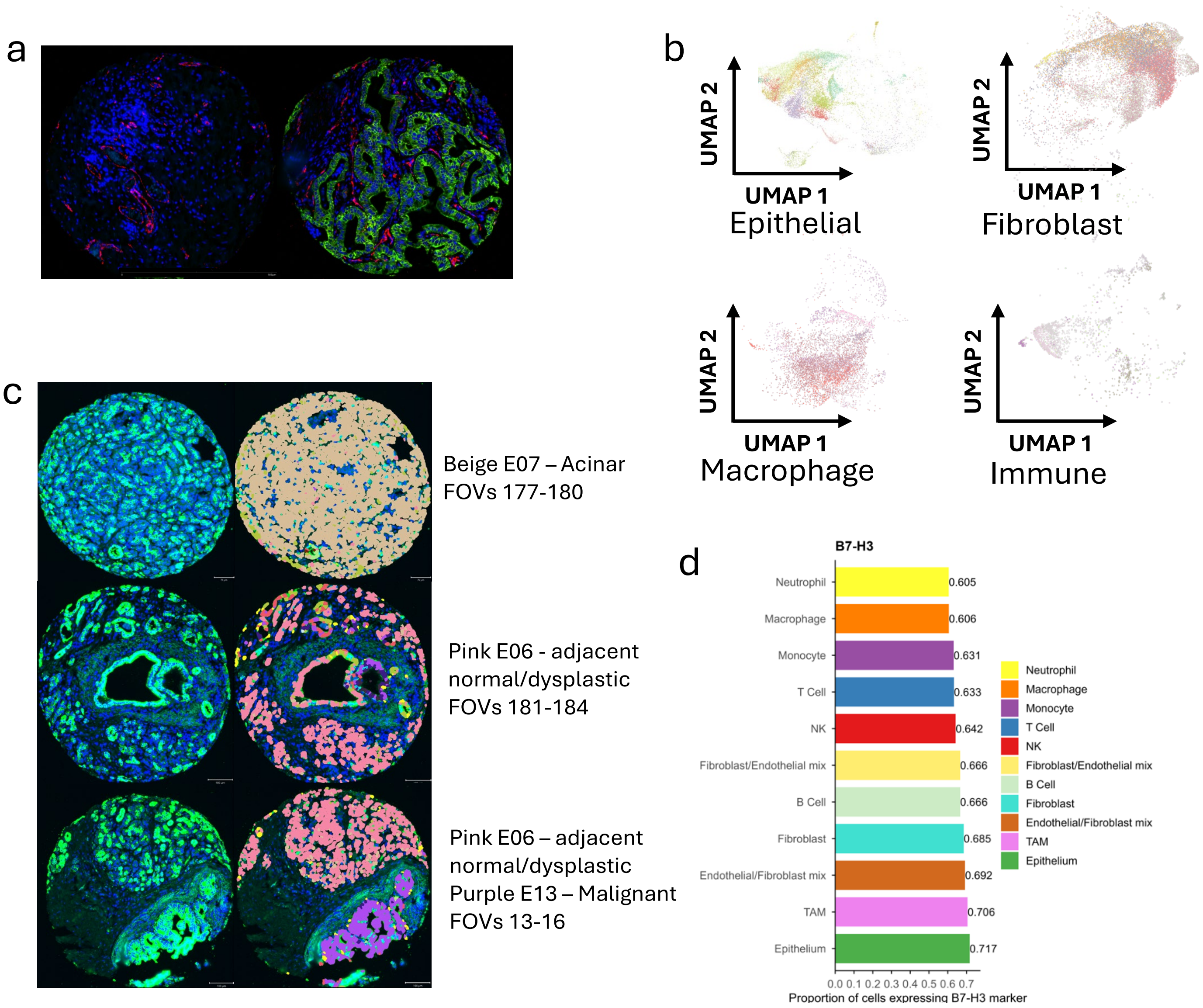


***Supplemental Figure 3:*** *Cell typing reveals B7-H3 enriched epithelial, fibroblast, and myeloid populations in naïve PDAC*

**a).** Multiplex immunofluorescent image core example showing overlap between CD31 marker expression. Red – CD31 (endothelial marker), Blue – Dapi, Green – PanCk. Image on left has clear endothelial structure as confirmed by pathologist, and image on right shows fibroblast staining morphology, intensity of staining is equal, resulting in poor separation between this mixed population, **b).** UMAPs showing subclusters of fibroblast (top left), macrophage/monocyte (bottom left) and immune (bottom right), **c).** Three example cores showing multiplex fluorescent staining (PanCk – green, Nuclei – Dapi), with matched epithelial subclusters overlaid, including presumed normal acinar (E07, top core), presumed adjacent normal mixed with dysplastic (E06, middle core), presumed adjacent normal mixed with dysplastic (E06) and malignant (E13), **d).** Bar chart showing proportion of cell expressing B7-H3 in parent cell types, ranging from 0-1, coloured by parent cell type.

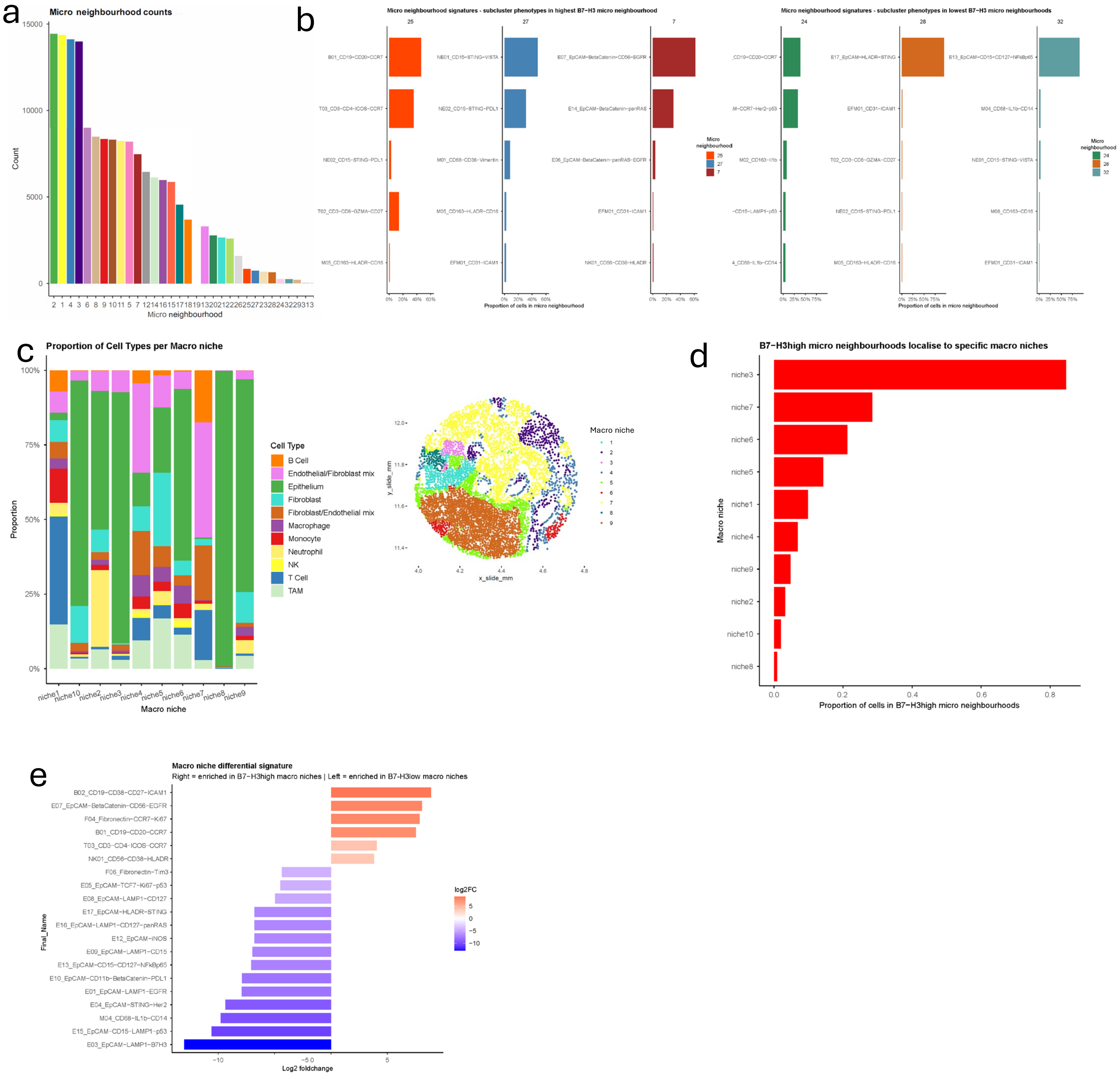


***Supplemental figure 4:*** *B7-H3 defines multiscale tumour states in PDAC, organising fibro-immune spatial neighbourhoods in B7-H3 high disease while preserving epithelial-centric architecture in B7-H3 low tumours*

**a).** Ranked bar chart showing the most common micro neighbourhoods across the entire treatment naïve pancreatic cancer validation cohort. **b).** Top 5 cell types associated with B7-H3 high and low ranked micro neighbourhoods and an example false image core with the micro neighbourhood displayed. **c).** Stacked bar chart showing proportion parent cell types of per macro niche. Macro niche were generated by local cell-type composition within a fixed physical radius. For each cell, neighbouring cells within a 50 µm radius were identified using a fast k-nearest neighbour search (RANN), for each focal cell, we computed a neighbourhood composition vector by counting the number of neighbouring cells assigned to each parent lineage (parent cell type), producing a per-cell composition matrix. Composition vectors were centred and scaled, denoised using principal component analysis (top 10 PCs), and clustered using k-means (k = 10; fixed random seed) to define macro-scale structural niches. **d).** Ranked bar chart showing the proportion of cells in B7-H3 high micro neighbourhoods (combined 25, 7, 27, 29, 5, and 9 networks) localised to all macro niches. **e).** Ranked bar chart showing top differentially expressed cell types in B7-H3 high (combined niches 3 and 7) and low (combined niches 8 and 10) macro niches

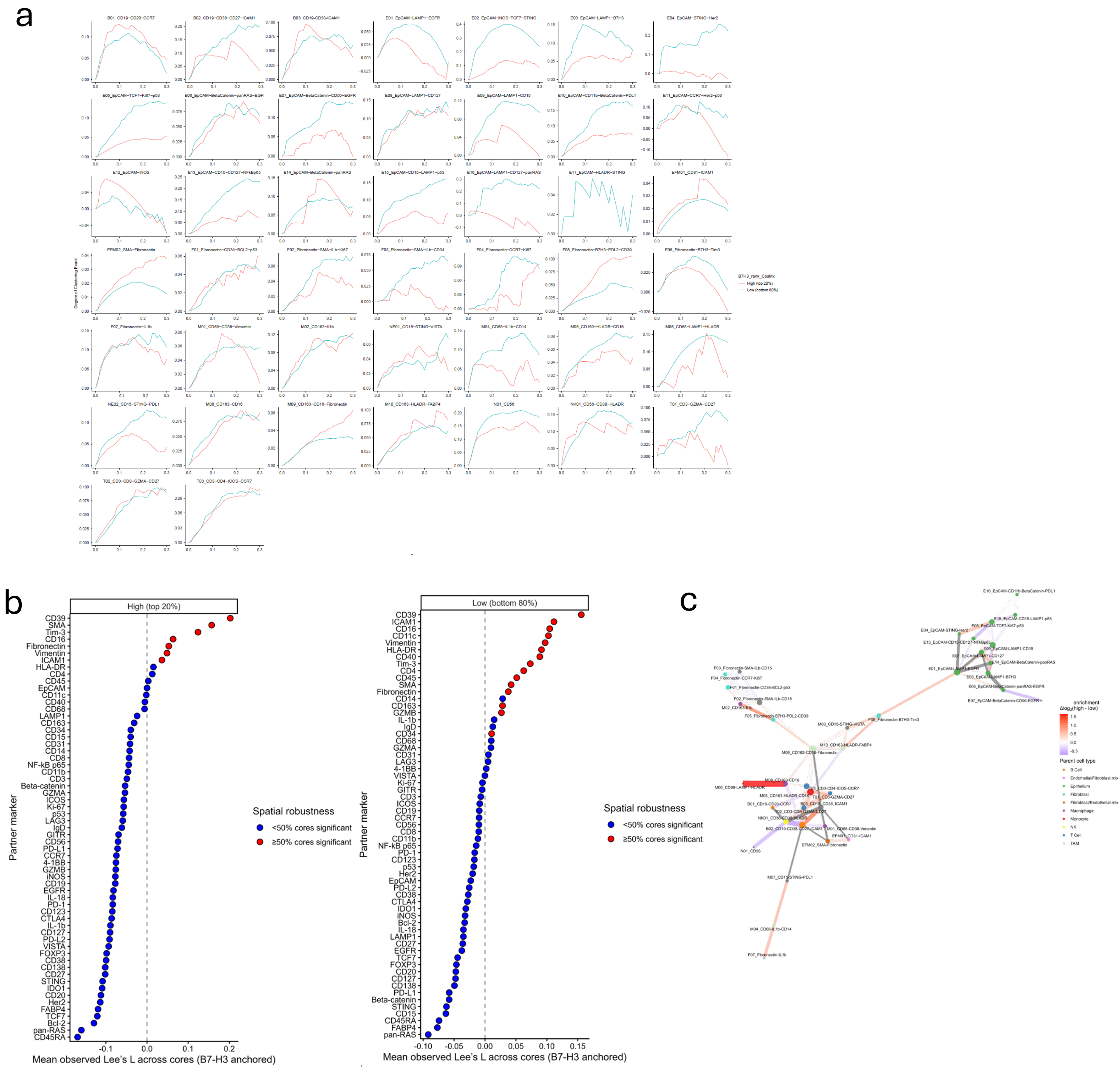


***Supplemental figure 5:*** *Spatial Dependency Analyses Reveal B7-H3 Centered Fibro-Immune Niches.*

**a).** Univariate Ripley's K curve for top phenotypes of interest showing Degree of clustering exact for full radius (mm). B7-H3 ranks are indicated by coloured lines (B7-H3high - red, B7-H3low – blue), **b).** Ranked dot plot showing mean observed Lee's L across cores (range = -0.1 to 0.2), with B7-H3 as the anchor and all other markers as the partner, split by ranked B7H3 patient group. Spatial robustness is displayed, showing cores with <50% significant coloured blue, and cores with >50% significant coloured red, **c).** Full cell to cell spatial interaction network plot for ranked B7-H3 expression. Edge thickness indicates $\log_2$ enrichment of cell-to-cell interactions calculated by Fisher's exact test, reflecting the strength of non-random spatial co-localisation. Node size indicates network degree (number of significantly enriched interaction partners), identifying highly connected hub cell types. Node colour represents parent cell type.

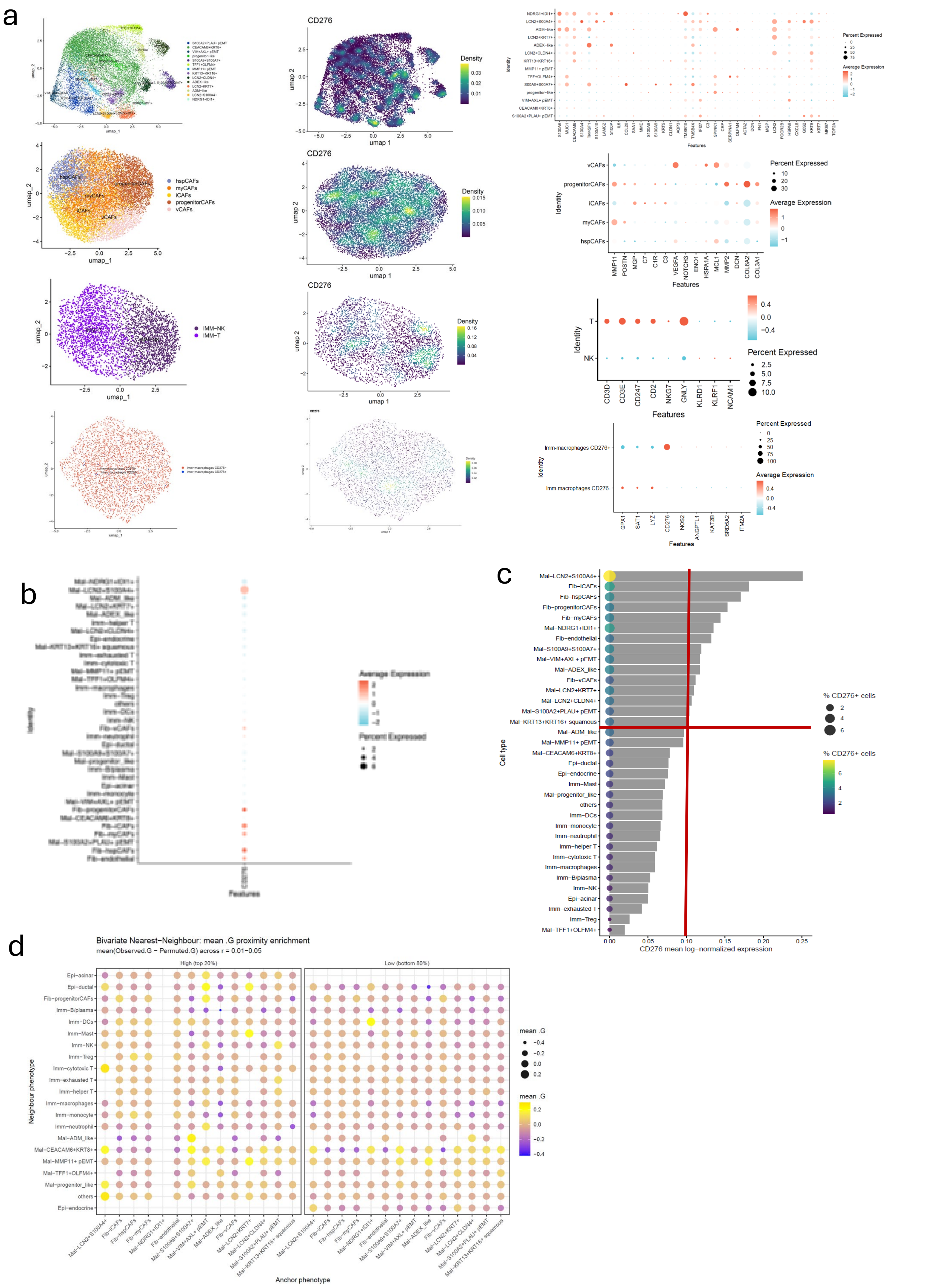


***Supplemental figure 6:** RNA panel reproduces CD276 (*B7-H3*) cell-type enrichment and spatial structure seen in protein*

**a).** UMAP representing subclusters generated by waterfall cell typing for epithelial, fibroblast and immune parent cell types (left), UMAP showing normalized *CD276 (*B7-H3*)* expression, and a dot plot showing differentially expressed markers, **b).** Dot plot showing average normalized B7-H3 expression across final subclustered cell types, **c).** Ranked bar chart showing mean log-normalized expression across final subclustered cell types. The size of the dot of the left indicates the percentage of *CD276 (*B7-H3*)* expressing cells. The cells found in the top right of the red line are the highest and largest *CD276 (*B7-H3*)* expressors, **d).** Dot plot showing Bivariate Nearest neighbour mean proximity enrichment between *CD276 (*B7-H3*)* ranked patients and all other fine-grained subclusters, coloured by cell types, size of dot indicates enrichment, radius investigated 10-50μm.
