## Supplemental Tables for "B7-H3 Orchestrates Fibro-Immune Spatial states in Pancreatic Cancer"

| **Discovery Naïve Cohort** | | | | | |
| --- | --- | --- | --- | --- | --- |
| **Characteristic** | ***n = 28*** | **Median DSS** | ***P (Log-Rank)*** | ***RFS*** | ***P (Log-Rank)*** |
|  |  | **(months)** |  |  |  |
| **Age** |  |  |  |  |  |
| <65 | 13 | 23.1 | 0.4 | 11.2 | 0.3 |
| >65 | 15 | 24.47 |  | 15.57 |  |
| Unknown | NA |  |  |  |  |
| **Sex** |  |  |  |  |  |
| M | 20 | 23.1 | 0.8 | 11.3 | 1 |
| F | 8 | 22.44 |  | 16.61 |  |
| Unknown | NA |  |  |  |  |
| **T Stage (AJCC 8^th^)** |  |  |  |  |  |
| T1 | 2 | 9.09 | 0.04 | 7.95 | 0.3 |
| T2 | 9 | 22.25 |  | 15 |  |
| T3 | 3 | 16.7 |  | 10 |  |
| T4 | NA |  |  |  |  |
| Unknown | 14 |  |  |  |  |
| **Lymph Node** |  |  |  |  |  |
| LN0 | 2 | 62.9 | 0.5 | 60.5 | 0.4 |
| LN1 | 11 | 18.55 |  | 14.65 |  |
| Unknown | 15 |  |  |  |  |
| **Resection Margin** |  |  |  |  |  |
| R0 | 24 | 16.45 | 0.3 | 13.45 | 0.2 |
| R1 | 4 | 73.5 |  | 62.9 |  |
| Unknown | 0 |  |  |  |  |
| **N Stage (AJCC 8^th^)** |  |  |  |  |  |
| N0 | 2 | NA | 0.07 | NA | 0.07 |
| N1 | 21 | 17 |  | 11.7 |  |
| N2 | NA |  |  |  |  |
| Unknown | 5 |  |  |  |  |
| **Grade / Tumour Differentiation** |  |  |  |  |  |
| I / Well | 3 | 36 | 0.2 | 18.9 | 0.3 |
| II / Moderate | 20 | 17.1 |  | 11.7 |  |
| III / Poor | 5 | 9.75 |  | 4.6 |  |
| IV/Undifferentiated | NA |  |  |  |  |
| Unknown | 0 |  |  |  |  |
| **Vascular Invasion** |  |  |  |  |  |
| Negative | 10 | 27.5 | 0.5 | 19.5 | 0.2 |
| Positive | 18 | 16.9 |  | 10.7 |  |
| Unknown | 0 |  |  |  |  |

**Supplemental Methods Table 1.** Treatment-naïve discovery PDAC cohort characteristics. Table showing treatment-naïve discovery PDAC cohort patients with clinical characteristics associated with disease specific survival (DSS) and recurrence free survival (RFS). Clinical features include T Stage (AJCC8th), resection margin status, lymph node status and vascular invasion.

| **Validation Naïve Cohort** | | | | | |
| --- | --- | --- | --- | --- | --- |
| **Characteristic** | ***n = 79*** | **Median DSS** | ***P (Log-Rank)*** | ***RFS*** | ***P (Log-Rank)*** |
|  |  | **(months)** |  |  |  |
| **Age** |  |  |  |  |  |
| <65 | 44 | 22.8 | 0.9 | 14.6 | 0.8 |
| >65 | 35 | 20 |  | 9.37 |  |
| Unknown | NA |  |  |  |  |
| **Sex** |  |  |  |  |  |
| M | 46 | 17.5 | 0.7 | 11.01 | 0.4 |
| F | 33 | 24 |  | 15.6 |  |
| Unknown | NA |  |  |  |  |
| **T Stage (AJCC 8^th^)** |  |  |  |  |  |
| T1 | 8 | 25.4 |  | 18.2 |  |
| T2 | 42 | 25 | 0.02 | 14.2 | 0.03 |
| T3 | 18 | 10.9 |  | 8.9 |  |
| T4 |  |  |  |  |  |
| Unknown | 11 |  |  |  |  |
| **Lymph Node** |  |  |  |  |  |
| LN0 | 10 | 36.5 | 0.04 | 25.9 | 0.03 |
| LN1 | 63 | 20 |  | 12.2 |  |
| Unknown | 6 |  |  |  |  |
| **Resection Margin** |  |  |  |  |  |
| R0 | 21 | 25.8 | 0.1 | 16.1 | 0.05 |
| R1 | 52 | 17.5 |  | 11 |  |
| Unknown | 6 |  |  |  |  |
| **N Stage (AJCC 8^th^)** |  |  |  |  |  |
| N0 | 8 | 23.1 | 0.5 | 14.8 | 0.8 |
| N1 | 60 | 21.2 |  | 12.2 |  |
| N2 | 0 |  |  |  |  |
| Unknown | 11 |  |  |  |  |
| **Grade / Tumour Differentiation** |  |  |  |  |  |
| I / Well | 4 | 24 |  | 11.6 |  |
| II / Moderate | 41 | 19.1 |  | 11 |  |
| III / Poor | 28 | 19.1 | 0.9 | 27.4 | 0.1 |
| IV/Undifferentiated | 0 |  |  |  |  |
| Unknown | 6 |  |  |  |  |
| **Vascular Invasion** |  |  |  |  |  |
| Negative | 39 | 24.5 | 0.2 | 15.1 | 0.2 |
| Positive | 34 | 16.3 |  | 10.2 |  |
| Unknown | 6 |  |  |  |  |

**Supplemental Methods Table 2.** Treatment-naïve validation PDAC cohort characteristics. Table showing treatment-naïve validation PDAC cohort patients with clinical characteristics associated with disease specific survival (DSS) and recurrence free survival (RFS). Clinical features include T Stage (AJCC8th), resection margin status, lymph node status and vascular invasion.

| **Marker** | **Company** | **Retrieval conditions** | **Antibody dilution** | **Secondary** |
| --- | --- | --- | --- | --- |
| CD3 | Agilent (A0452) | High pH TRS buffer (K8004, Agilent) | 1in50 | EnVision rabbit (K4003, Agilent) |
| CD8 | Leica (NCL-L-CD8-4B11) | ER2 20 mins | 1in75 | EnVision mouse (K4001, Agilent) |
| CD68 | Agilent (M0876) | ER2 20 mins | 1in500 | EnVision mouse (K4001, Agilent) |
| CD163 | Leica (NCL-L-CD163) | ER2 20 mins | 1in300 | EnVision mouse (K4001, Agilent) |

**Supplemental Methods Table 4.** IHC conditions for antibody of interests. Description of antibody marker (CD3, CD8, CD68, CD163), retrieval conditions, antibody dilution.

| **Compartment** | **Percentile** | **Cut point** | **High / Low (n)** | **P value** | **FDR** | **Chi-squared** | **Note** |
| --- | --- | --- | --- | --- | --- | --- | --- |
| Tumour | 50th | 8.1 | 14 / 11 (56% / 44%) | 0.003 | 0.021 | 8.65 | Significant - but biologically non-specific |
| Tumour | 55th | 8.3 | 12 / 13 (48% / 52%) | 0.005 | 0.021 | 7.74 | Significant - majority classified as 'High' |
| Tumour | 60th | 8.34 | 10 / 15 (40% / 60%) | 0.034 | 0.051 | 4.51 | Non-significant |
| Tumour | 65th | 8.4 | 10 / 15 (40% / 60%) | 0.034 | 0.051 | 4.51 | Non-significant |
| Tumour | 70th | 8.48 | 8 / 17 (32% / 68%) | 0.068 | 0.073 | 3.33 | Non-significant |
| Tumour | 75th | 8.6 | 7 / 18 (28% / 72%) | 0.007 | 0.022 | 7.22 | Significant - but not the most extreme expressors |
| Tumour | 80th | 8.62 | 5 / 20 (20% / 80%) | 0.018 | 0.04 | 5.62 | Selected cut point - biologically principled |
| Tumour | 85th | 8.78 | 4 / 21 (16% / 84%) | 0.042 | 0.053 | 4.15 | Non-significant |
| Tumour | 90th | 9.02 | 3 / 22 (12% / 88%) | 0.073 | 0.073 | 3.22 | Non-significant - n too small |
| Tumour | 95th | 9.42 | < 3 | NA | NA | NA | Insufficient n for testing |

**Supplemental Methods Table 5.** Log-rank test results for B7-H3 dichotomisation at each percentile threshold from the 50th to the 95th percentile within tumour compartments. n High and n Low refer to the number of patients in each group. The selected 80th percentile cut point is highlighted. Thresholds with fewer than 3 events per group were excluded (indicated by NA). Benjamin-Hochberg adjustment performed and Chi-squared statistic reported with 1 degree of freedom.
